# Rare Subset of T Cells Form Heterotypic Clusters with Circulating Tumor Cells to Foster Cancer Metastasis

**DOI:** 10.1101/2025.04.01.646421

**Authors:** David Scholten, Lamiaa El-Shennawy, Yuzhi Jia, Youbin Zhang, Elizabeth Hyun, Carolina Reduzzi, Andrew D. Hoffmann, Hannah F. Almubarak, Fangjia Tong, Nurmaa Dashzeveg, Yuanfei Sun, Joshua R. Squires, Janice Lu, Leonidas C. Platanias, Clive H Wasserfall, William J. Gradishar, Massimo Cristofanilli, Deyu Fang, Huiping Liu

## Abstract

The immune ecosystem is central to maintaining effective defensive responses. However, how immune cells in the periphery blood interact with circulating tumor cells (CTCs) – seeds of metastasis - remains largely understudied. Here, our analysis of the blood specimens (N=1,529) from patients with advanced breast cancer revealed that over 75% of the CTC-positive blood specimens contained heterotypic CTC clusters with CD45^+^ white blood cells (WBCs). Detection of CTC-WBC clusters correlates with breast cancer subtypes (triple negative and luminal B), racial groups (Black), and decreased survival rates. Flow cytometry and ImageStream analyses revealed diverse WBC composition of heterotypic CTC-WBC clusters, including overrepresented T cells and underrepresented neutrophils. Most strikingly, a rare subset of CD4 and CD8 double positive T (DPT) cells showed an up to 140-fold enrichment in the CTC clusters *versus* its frequency in WBCs. DPT cells shared part of the profiles with CD4^+^ T cells and others with CD8^+^ T cells but exhibited unique features of T cell exhaustion and immune suppression with higher expression of TIM-3 and PD-1. Single-cell RNA sequencing and genetic perturbation studies further pinpointed the integrin VLA4 (α4β1) in DPT cells and its ligand VCAM1 in tumor cells as essential mediators of heterotypic WBC-CTC clusters. Neoadjuvant administration of anti-α4 (VLA4) neutralizing antibodies markedly blocked CTC–DPT cell clustering and inhibited metastasis for extended survival in preclinical mouse models *in vivo*. These findings uncover a pivotal role of rare DPT cells with immune suppressive features in fostering cancer dissemination through direct interactive clustering with CTCs. It lays a foundation for developing innovative biomarkers and therapeutic strategies to prevent and target cancer metastasis, ultimately benefiting cancer care.

**Brief summary:** Our findings uncover a fostering role of immune-suppressive T cells in contact with circulating tumor cells and identify therapeutic approaches to eliminate devastating cancer metastasis.

**Graphical Abstract:** 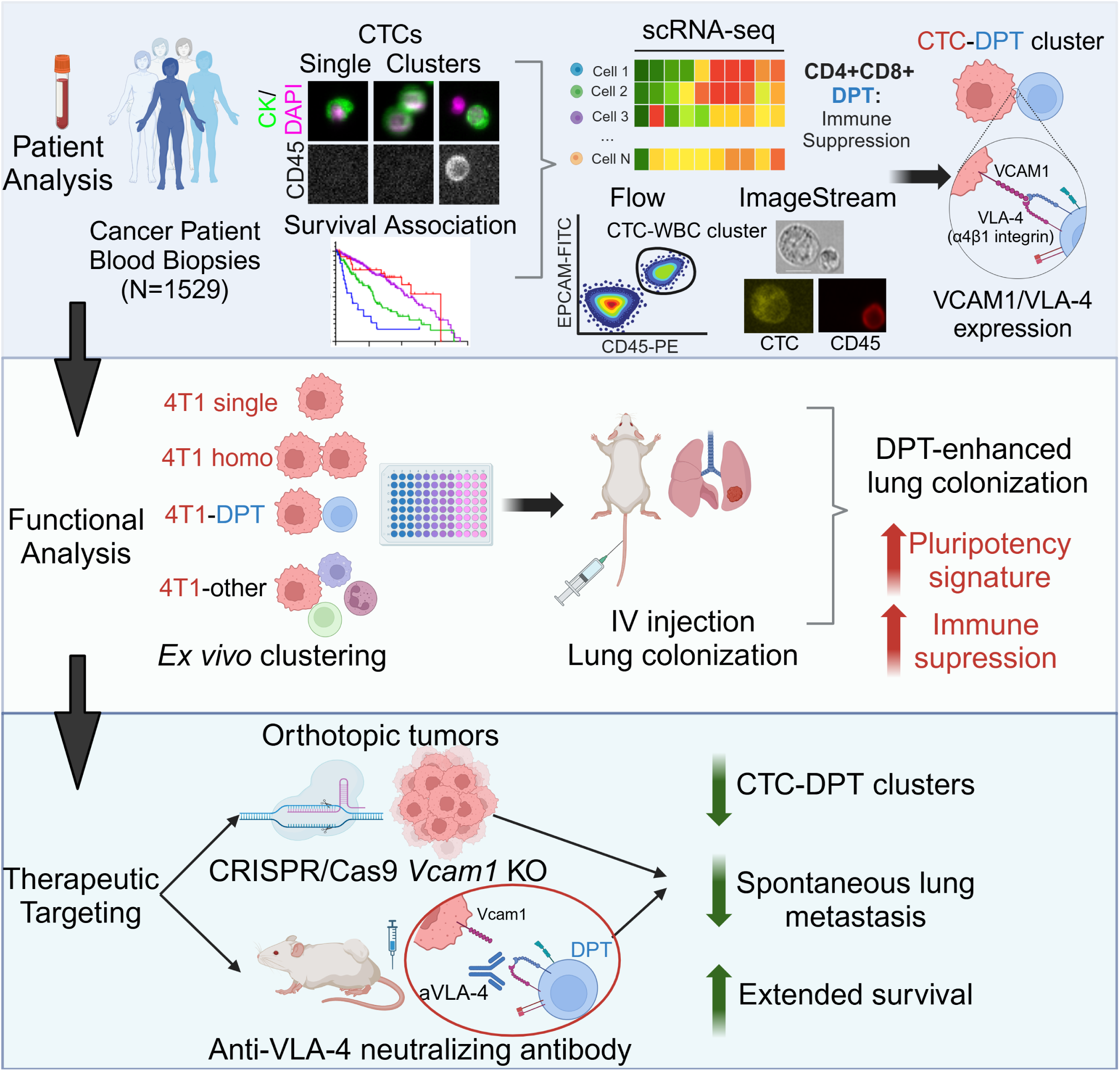

## Introduction

Circulating tumor cells (CTCs), invasive tumor cells that have entered the bloodstream with intrinsic regenerative properties and adaptive plasticity, have long been recognized as metastatic seeds of many cancers ^1–13^. The traditional paradigm of single cell-mediated metastasis has been reshaped by recent discoveries that clusters of multiple tumor cells (homotypic CTC clusters) show enhanced stemness and up to 50-fold greater metastatic potential than single CTCs, correlating with worse outcomes ^6,14–21^. While tumor cell fitness plays a critical role in CTC dissemination, it is pivotal - albeit technically challenging - to elucidate how immune cells, i.e. white blood cells (WBCs), engage with CTCs and influence metastasis.

The development and implementation of various technologies have greatly facilitated the clinical characterization of CTCs ^21–25^. Among those, an FDA-cleared CellSearch platform with EpCAM enrichment and immunofluorescence staining has been widely used in detecting CTCs (DAPI^+^ Cytokeratin/CK^+^ CD45^−^) from patients with epithelial cancers, such as breast^5^, prostate^26^ and colon ^27^.

Over the last 10 years, we have employed CellSearch for CTC detection in blood samples from patients with stage III and IV breast cancer (N=1,529). Notably, heterotypic CTC-WBC clusters were more frequently detected than homotypic CTC clusters in CTC-positive patients, especially those with triple-negative breast cancer and luminal B, and prognostically associated with poor survival, which is exacerbated in Black patients. We then examined the cellular composition and molecular characteristics of heterotypic CTC-WBC clusters. Establishing complementary approaches of flow cytometry and ImageStream cytometry for CTC analysis ^17,28^, we identified multiple immune cell types within heterotypic CTC clusters, including the relatively enriched T cells, under-represented neutrophils, and other less frequently detected cells. Most surprisingly, a rare subset of T cells in circulation, CD4 and CD8 double-positive T (DPT) cells, showed the highest enrichment (140-fold) relative to WBCs, thus forming the heterotypic clusters.

DPT cells ^29^ are extremely rare in the periphery blood even though they are bona fide, mature T cells during selective development ^30–32^. To our knowledge, the role of DPT cells in cancer metastasis or CTC clusters has never been reported. We set out to determine the functional contribution of heterotypic DPT-CTC clusters to metastatic seeding and to elucidate the molecular mechanisms underlying DPT-CTC clustering.

## Results

### Clinical associations of single and clustered CTCs in breast cancer

At the Northwestern Circulating Tumor Cell Core, we established a clinical protocol of blood CTC tests for patients with stage III and IV breast cancer from 2016 to 2025 (N=1,529) using the CellSearch platform coupled with image scans (**Figure 1A, Extended Table S1**). In addition to the enumeration report of single CTCs (DAPI^+^ CK^+^ CD45^−^), we manually reviewed the immunofluorescence images of EpCAM bead-precipitated cells for the identification of homotypic CTC clusters (>=2 CTCs/cluster) and heterotypic CTC clusters with CD45^+^DAPI^+^ WBCs (**Figure 1A-B, S1**). Of these tests, 43.23% were CTC positive (>=5 single CTCs within 7.5 mL blood). Of CTC-positive biospecimens (N=661), 44.02% contained homotypic clusters (at least 1 cluster), whereas 75.49% were positive for heterotypic clusters (at least 1 cluster) (**Figure 1B**). The range, mean, and median counts are 0-17,427; 47; 0 for single CTCs; 0-2,265; 3; 0 for homotypic clusters; and 0-1,017; 6; 2 for heterotypic clusters with a higher detection frequency than homotypic clusters but lower than single CTCs (**Figure 1C**, P<0.0001 Wilcoxon matched-pairs signed rank test). Most heterotypic CTC-WBC clusters contained only 1 immune cell with 1 tumor cell (**Figure S1B**).

**Figure 1.**
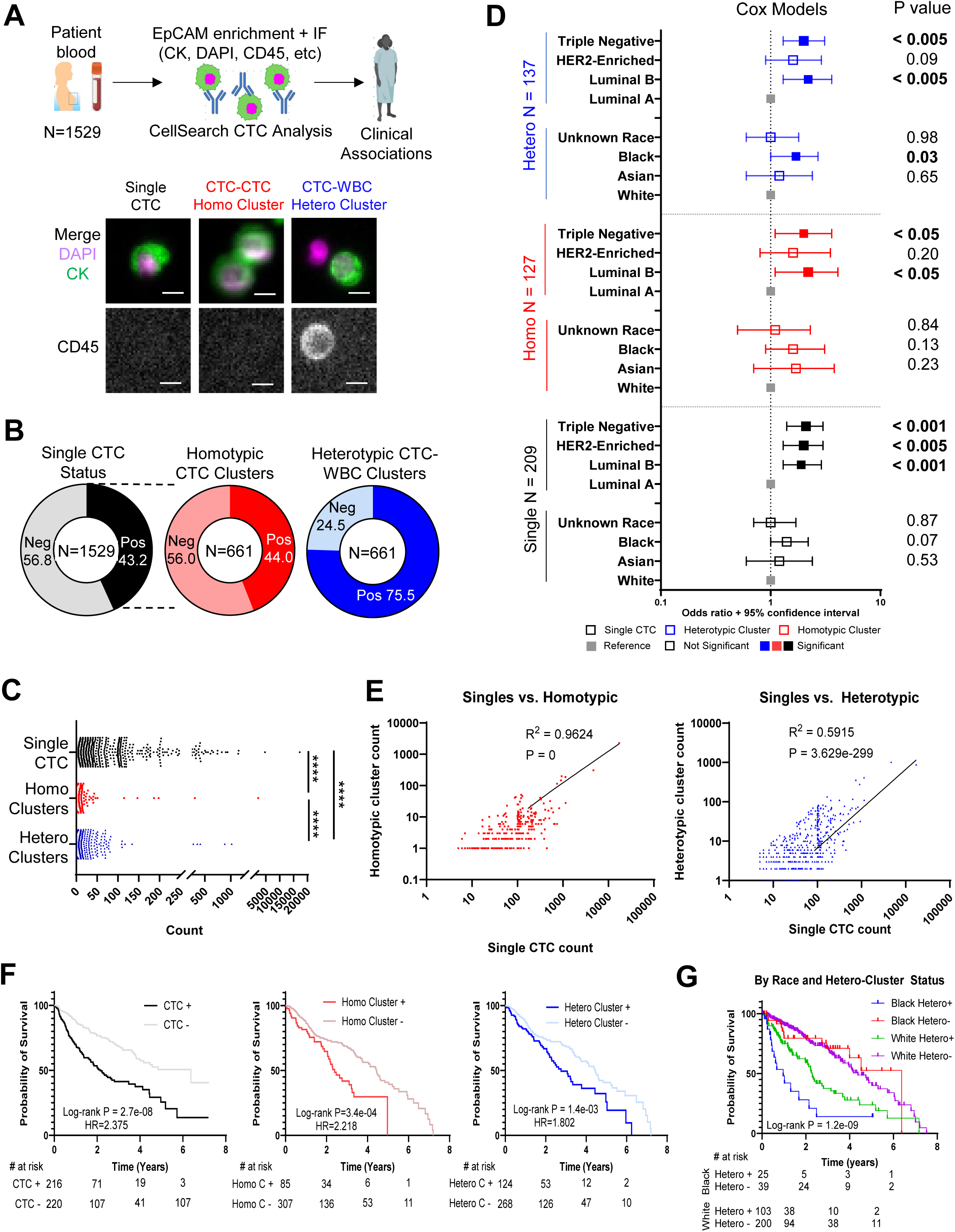
CTC frequencies in the blood biopsies of breast cancer patients and their clinical associations. **A.** Top panels: Schematic of CTC analysis via CellSearch (EpCAM bead pulldown and subsequent immunofluorescence staining for image scanning) with blood specimens drawn from patients with breast cancer (N=1529). Bottom panels: Representative CellSearch images of single CTC (left), homotypic CTC-CTC cluster (middle), or heterotypic CTC-WBC cluster (right) with merge channels of cytokeratin (CK) (green) and DAPI (magenta) as well as a single CD45 channel (Scale bar = 10 μm). **B.** Frequency of CellSearch-detected CTC positive tests/scans (≥ 5 CTCs within 7.5 mL blood) among breast cancer patient biopsies (left, N=1,529), and frequencies of homotypic CTC-CTC clusters (middle) and heterotypic (right) CTC-WBC clusters among CTC^+^ biospecimens (N=661). **C.** Counts of single CTCs and homotypic and heterotypic CTC clusters per 7.5 mL blood in 1,529 CellSearch tests. The range, mean, and median are 0-17,427; 47; 0 for single CTCs; 0-2,265; 3; 0 for homotypic clusters; and 0-1,017; 6; 2 for heterotypic clusters. P<0.0001 for any two-group comparison using the Wilcoxon matched-pairs signed rank test. **D.** Cox proportional hazard model odds ratio plot with 95% confidence interval for risk of single CTCs (black), homotypic CTC-CTC clusters (red), and heterotypic CTC-WBC clusters (blue) among subtypes of breast cancer and self-identified racial groups of the patients. Filled squares highlight significant features calculated using the Wald test (P<0.05). **E.** Scatter plots of single vs. homotypic clusters and single vs. heterotypic clusters with Pearson correlation coefficient and two-tailed p-value. **F.** Kaplan-Meier survival curves of patients positive for single CTCs (≥5), homotypic clusters (≥1), or heterotypic clusters (≥1) versus the patients with negative results. Log-rank (Mantel-Cox) test p-values and hazard ratio (HR) are displayed. **G.** Kaplan-Meier survival curves of patients with breast cancer, divided by race (Black and White) and heterotypic cluster status. Log-rank (Mantel-Cox) test p-value shown.

We performed a Cox proportional hazard analysis to determine which patient variables would be risk factors for single CTCs, homotypic clusters, or heterotypic clusters (**Figure 1D**). Compared to the White group, self-identified Black or African American patients had a higher risk specifically for heterotypic clusters with no significant difference for single CTCs nor homotypic clusters. When luminal A breast cancer subtype served as the reference control, luminal B, HER2-enriched, and triple negative breast cancer (TNBC) showed higher risks of single CTCs. In contrast, only TNBC and luminal B had higher risks for homotypic and heterotypic clusters.

Single CTC counts positively correlate with both cluster types, in a stronger correlation with homotypic cluster counts (R^2^ = 0.9624) than heterotypic cluster count (R^2^ = 0.5915) (**Figure 1E**), implying that additional factors of the WBCs might contribute to heterotypic cluster formation. Based on the clinical follow-up data, the Kaplan-Meier analysis of these patients demonstrated new prognostic values of heterotypic CTC-WBC clusters and confirmed those of single CTCs ^5^ and homotypic CTC clusters ^6,17^, all of which correlate with unfavorable survival (**Figure 1F**). Notably, when stratified by race and the status of heterotypic clusters, Black patients positive for heterotypic clusters had the shortest overall survival versus the other three groups (**Figure 1F**). These data provide novel insights into the clinical utility of CTCs, especially heterotypic CTC-WBC clusters associated with breast cancer outcomes.

### DPT cells are enriched in heterotypic CTC-WBC clusters and promote seeding

We continued to examine the WBC composition in the heterotypic clusters and their functional contribution to CTC-mediated seeding. Because CellSearch is limited to four-channel immunofluorescence staining with only one open for customized marker analysis, we utilized established flow cytometry ^15,33^ and imaging flow approaches (ImageStream and BD CellView) to characterize WBCs in heterotypic CTC clusters, based on expanded channels and cellular images (**Figure 2A-B**). After red blood cell lysis, we profiled the blood cells of 26 patients with breast cancer using extensive markers ^34,35^ (plus forward and side scatter channels) to identify T cells, B cells, NK cells, monocytes, and neutrophils in single WBCs and heterotypic clusters (**Figure 2B, S2**). Compared to single WBCs in circulation, heterotypic CTC clusters included over-represented T cells (32.6%) and NK (12.6%), underrepresented neutrophils (30.2%), and monocytes (19.9%) and B cells (4.7%) without significant changes (**Figure 2C-E**). The most striking identification was CD4^+^CD8^+^ DPT cells (14.2%) with a 140-fold enrichment in the clusters versus its rare presence in WBCs (0.1%), in which double positivity was confirmed *via* ImageStream imaging cytometry (**Figure 2C-E**). Instead, the single positive CD8 and CD4 T cells made 13.2% and 5.6% of the heterotypic clusters, respectively (**Figure 2C-E**).

**Figure 2.**
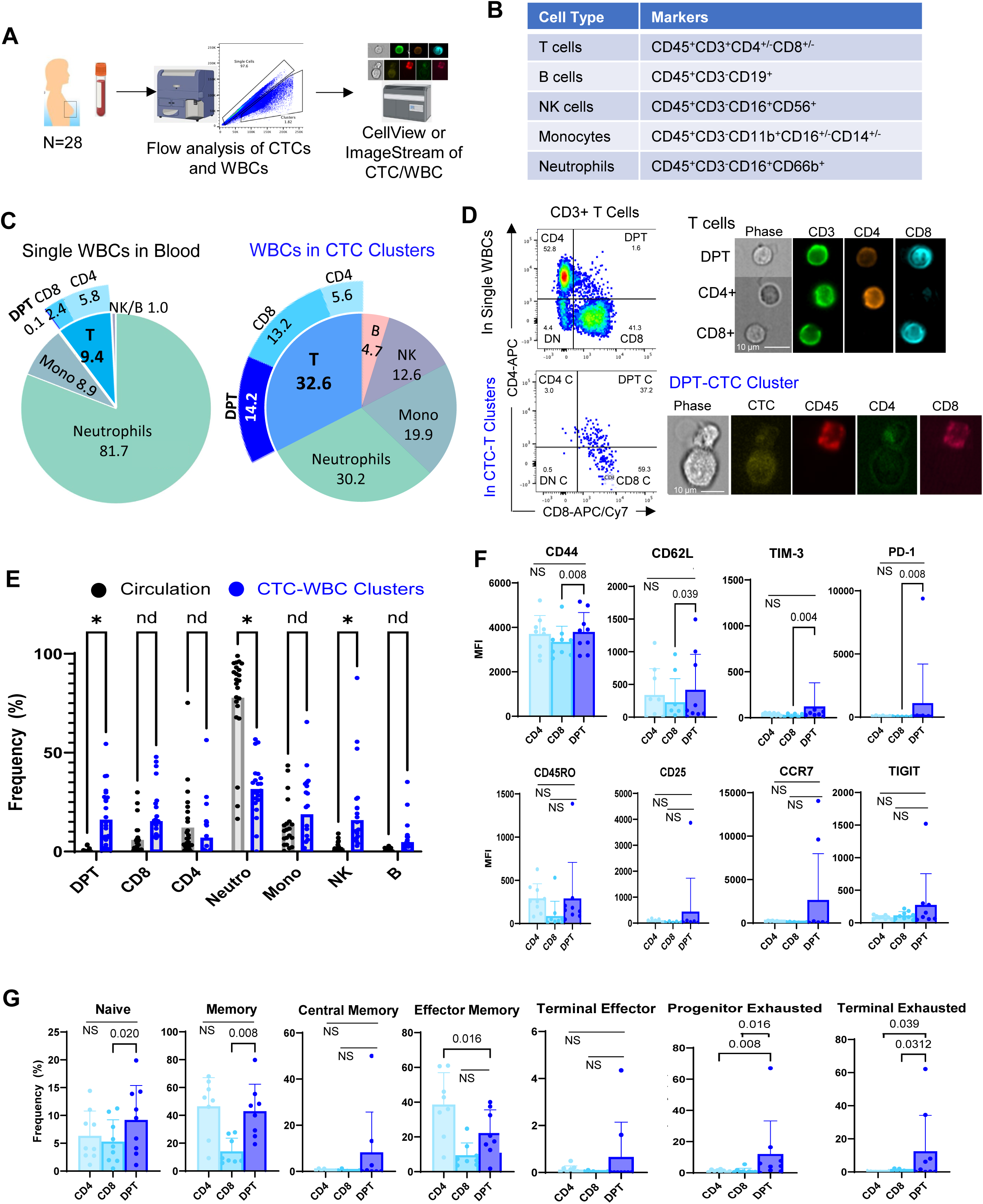
Double positive T (DPT) cells are 140-fold enriched in CTC-WBC clusters compared to single WBCs. A. Schematic of analyzing the frequency of broad classes of immune cell compositions in the blood biopsies (single WBCs and CTC-WBC clusters) of breast cancer patients via flow cytometry and ImageStream/CellView. N = 26 patients (n=1,402 CTC-WBC clusters). B. List of representative WBCs with surface markers used to identify immune cell types present in CTC-WBC clusters. C. Frequency of immune cells (neutrophils, monocytes, NK, and B) and different T cell populations (DPT, CD4+T, CD8+T) in patient blood (WBCs) (left pie charts) and CTC-WBC clusters (right pie charts). N = 26 patients (n=698 CTC-T cell clusters). D. Left panels: flow dot plots of CD3+ T cells in single WBCs (top) and heterotypic CTC-WBC clusters (bottom). Top right panels: ImageStream photos of CD8+CD4+ double positive T (DPT), CD4+T, and CD8+T cells. Bottom right panels: Representative images of CTC-DPT cluster via ImageStream imaging cytometry. E. Frequency of subset T, neutrophils, monocytes, NK, and B from individual patients. Multiple paired t test, *, q < 0.05. N = 26 F. Median fluorescence intensity (MFI) of various T cell markers (CD44, CD62L, CD45RO, CCR7, TIM-2, PD-1, CD25, TIGIT) in human DPT cells of breast cancer patients compared to CD4^+^ and CD8^+^ single positive T cells, detected by flow cytometry. A Wilcoxon matched-pairs signed-rank test with significant p-value changes displayed, NS=not significant. N = 8. G. Phenotypic characterization of human DPT cells in breast cancer patients compared to CD4^+^ and CD8^+^ single positive T cells, including naïve (CD62L^+^CD44^−^), memory (CD45RO^+^), central memory (CD45RO^+^CCR7^+^), effector memory(CD45RO^+^CCR7^−^), terminal effector (PD-1^+^TIM3^−^), progenitor exhausted (TIM3^+^PD-1^−^), terminal exhausted cells (PD-1^+^TIM3^+^). N = 8 patients. Wilcoxon matched-pairs signed rank test between DPT and CD4/CD8, P <0.05 shown; NS=not significant. N = 8 breast cancer patients.

While thymocytes could go through a double positive stage in T cell development and could be fully mature T cells ^29^, we found that DPT cells were relatively rare in the periphery in the blood analyses (shown above). In collaboration with the HuBMAP consortium ^36,37^, we obtained the human spleen images of spatial immunofluorescence staining and identified rare DPT cells close to the germinal centers (**Figure S3**). We further characterized human DPT cell features in the blood biopsies of breast cancer patients using flow cytometry (**Figure S4A**). DPT cells displayed a marker profile closely resembling that of CD4^+^ T cells. However, compared to CD8^+^ T cells, they had slightly higher expression levels of CD44 (activation), CD62L (trafficking, also known as L-Selectin), and exhaustion (TIM3 and PD-1) (**Figure 2F**). In line with this, DPT cells had a similar distribution as CD4^+^ T cells but portrayed higher frequencies than CD8+ T cells in naïve (CD62L^+^CD44^−^), memory (CD45RO^+^), and terminal effector (PD-1^+^TIM3^−^) subsets. One notable exception was a slightly lower frequency of effector memory (CD45RO^+^CCR7^−^) in DPT cells than CD4^+^ T cells (**Figure 2G**). Nevertheless, DPT uniquely showed higher frequencies of both progenitor-exhausted (TIM3^+^PD-1^−^) and terminal-exhausted (PD-1^+^TIM3^+^) phenotypes relative to single positive T cells (**Figure 2G**). This suggests that DPTs in cancer patients have an activated, exhausted, and immunosuppressive phenotype.

Next, we sought to determine the role of DPT cells in heterotypic cluster-mediated metastatic seeding during experimental lung colonization in syngeneic mouse models (**Figure 3A**). After splenocytes were harvested from Balb/c mice, DPT cells were isolated via high-speed sorting (see methods). These cells were pre-mixed at 4:1 with murine TNBC 4T1 cells (Luc2-tdTomato/L2T labeled ^38^) and seeded to a poly-HEMA coated 96 well plate for 6-hour cluster formation *ex vivo* (**Figure S5A**). Mixed cells and clusters were collected gently and immediately infused into the tail veins of Balb/c recipient mice.

**Figure 3.**
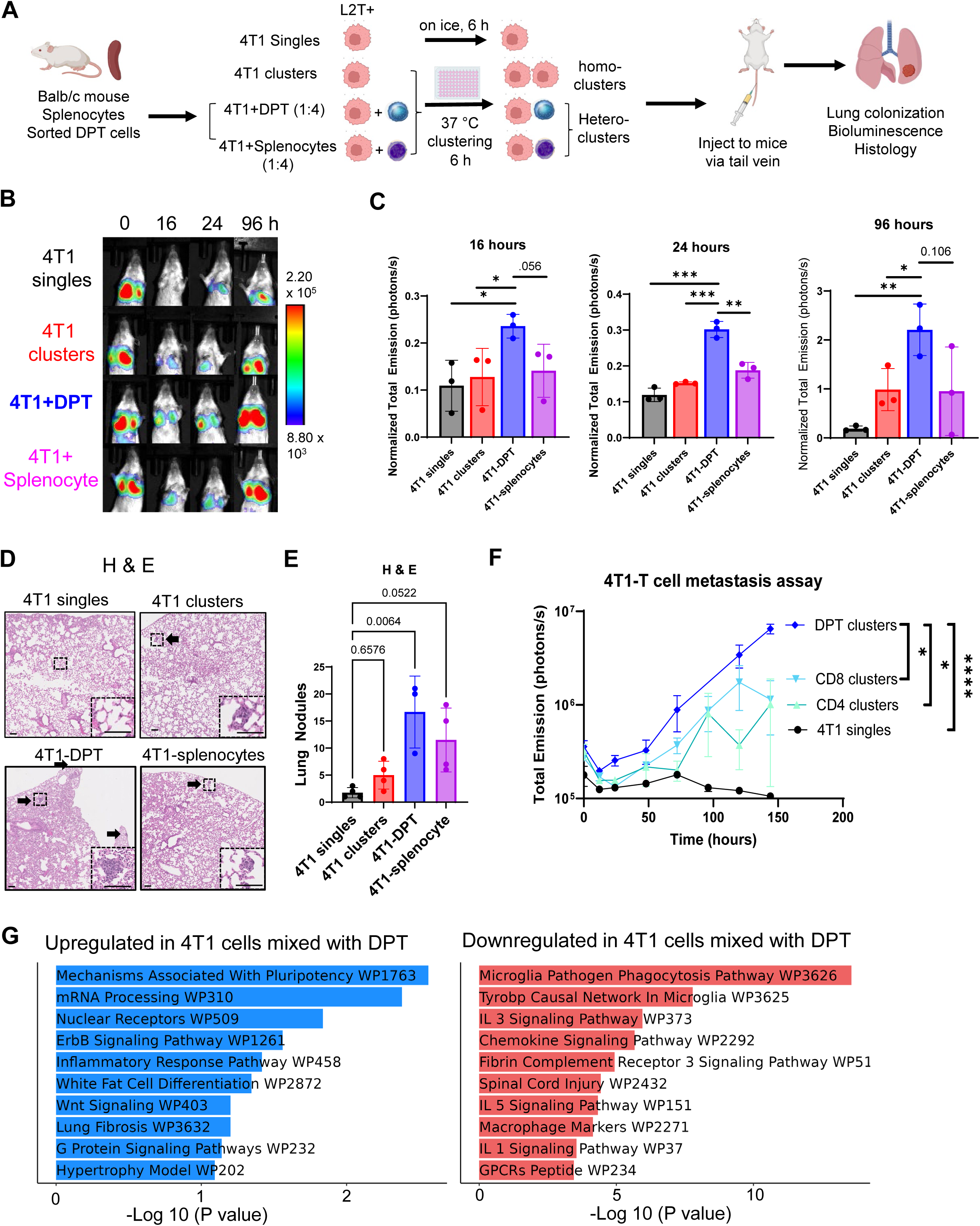
DPT-tumor cell clustering promotes metastasis formation in an experimental metastasis assay *in vivo*. **A.** Schematic depicting experimental design of mouse DPT isolation, clustering with L2T-labeled 4T1 tumor cells *ex vivo* (controls groups including 4T1 cells (singles and clusters), 4T1 and mouse splenocytes), tail vein infusion, and lung colonization monitored via bioluminescence imaging of L2T^+^ 4T1 cells and histology validation. **B-C.** Representative images (B) and quantification (C) of *in vivo* bioluminescent signals in mouse lungs of L2T^+^ 4T1 tumor cells after clustering and tail vein injection. Two-tailed *t*-test, n = 3 mice per group. **D-E.** Hematoxylin & eosin staining images with inserted regions of control or micrometastasis in the mouse lungs (D) and quantification of metastasis lesions of lung sections (E) on Day 6 after infusion of 4 groups of cells: 4T1 singles, 4T1 homo-clusters, 4T1-DPT clusters, and 4T1-splenocytes. Arrows pointing to metastatic lesions. Scale bars = 100 µm. Three-way ANOVA is used for P-value calculations. **F.** Repeated experiment of DPT-4T1 clustering-promoted metastatic seeding and colonization with single CD4+ and CD8+ T cell controls in mix-clustering with 4T1 cells. The graph represents mean ± standard error of the mean. Unpaired two tailed T test, *P<0.05, and ***P<0.001. N = 3 mice for each group. **G.** Enriched pathways upregulated (left) or downregulated (right) in 4T1 cells after being incubated with DPT cells vs. those with splenocytes for 6 hours, identified via Enrichr database (https://maayanlab.cloud/Enrichr/) (WikiPathways or WP) based on single-cell RNA sequencing (10x Genomics).data. The respective UMAPs for each group provided in Extend Figure 5B.

Compared to three control groups of 4T1 singles, 4T1 mixture with homotypic clusters, and 4T1-splenocyte mixture, 4T1-DPT mixture seeded to the lungs with the highest efficiency and strongest bioluminescence signal at 16, 24, and 96 hours (**Figure 3B, C**). H & E staining of the mouse lungs at 6 days post-injection validated the better colonization by 4T1-DPT clusters compared to other groups (**Figure 3D-E**).

As single positive T cells were part of the WBCs clustered with CTCs in patient blood, we also compared the seeding efficiency of the DPT-4T1 mixture with CD4-4T1 and CD8-4T1 mixtures. Similarly, infusion of DPT-4T1 cells resulted in the highest lung bioluminescent signal of disseminated tumor cells at 6 days post-tail vein injection compared to these two groups and other controls (**Figure 3F**). These data demonstrate that DPT cells of heterotypic clusters promote CTC-mediated pulmonary seeding and colonization.

To elucidate DPT interaction-mediated effects on tumors, we conducted scRNA-seq of the 4T1 tumor cells incubated with sorted DPTs and splenocytes for 6 hours of clustering. The tumor cells post DPT co-incubation partially overlapped the UMAP profiles with splenocyte-interacted tumor cells both differentially expressed genes (**Figure S5B, Extended Table S2**). Using the pathway analysis tool Enrichr ^39,40^, we identified DPT-upregulated pathways in tumor cells (pluripotency, mRNA processing, nuclear receptors, ErbB signaling, etc) as well as more significantly downregulated pathways in immune activation-related phagocytosis, IL-3, IL-5, and IL-1 signaling, chemokine signaling, etc (**Figure 3G, Extended Table 3**). These data suggest that the DPT interaction promotes stemness-related pluripotency and immune evasion of tumor cells.

### ITGB1/ITGA4 as drivers of heterotypic T cell clustering with tumor cells

While one previous study reported intratumoral DPT cells^41^, circulating DPT cells were rarely studied compared to other WBCs. To bridge this gap, we performed scRNA-seq of human peripheral WBCs from 19 breast cancer patients and 12 healthy controls (**Figure 4A**). After performing quality control and dataset integration, 8 broad immune cell types were identified in both cohorts (**Figure 4A, S6A-S6B**), including over-represented T, B, and NK cells in healthy controls along with more abundant intermediate monocytes in breast cancer patients (**Figure 4C**, top). However, when the patients were stratified by their CTC status into three groups, WBCs from CTC-positive patients (>= 5 CTCs) showed a higher proportion of T cells and classical monocytes than the other two cancer groups with low (1-5 CTCs) or no CTCs (**Figure 4C**, bottom, **Figure S6C**).

**Figure 4.**
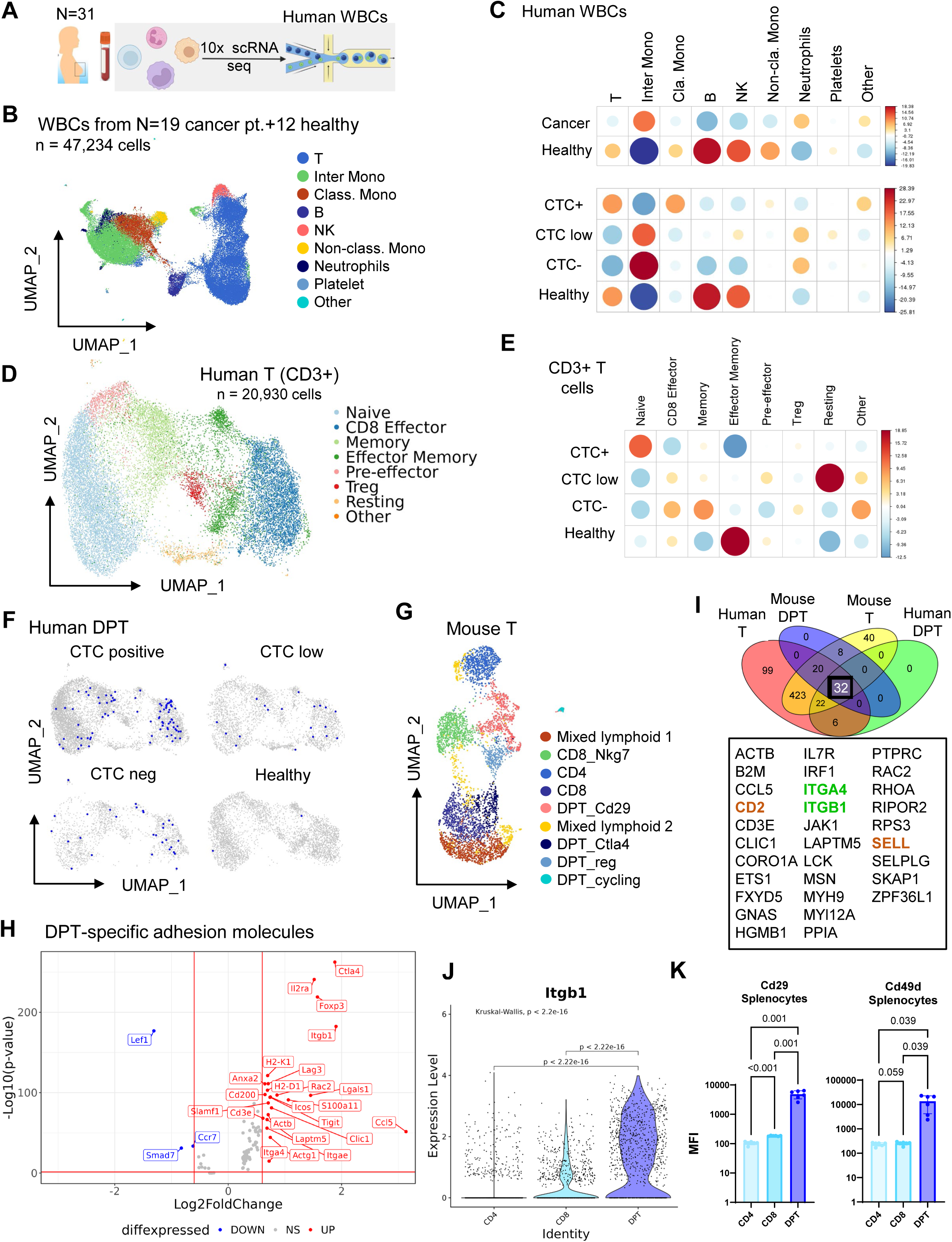
scRNA sequencing reveals enrichment of rare T cell subsets (DPT) in the blood of breast cancer patients, dependent on CTC status. A. Schematic depicting the isolation of human white blood cells (T cells via FACS) and subsequent 10X Genomics single-cell RNA sequencing. B. UMAP plots of 47,234 single white blood cells (WBCs) from 19 breast cancer patients (n=35,401 cells) and 12 healthy control liquid biopsies (n=11,833 cells), with broad immune cell subsets annotated. C. Correlation plots depicting chi-squared test residuals to determine over- or under-enrichment of each immune cell subset from the WBCs of breast cancer patients vs. healthy controls (top panel), and by CTC status (bottom panel), including breast cancer patients with CTC positive (≥5 CTCs), CTC low (1-4 CTCs), and CTC negative (0 CTCs) per 7.5 mL blood, and healthy controls (non-cancer). Dot size corresponds to the absolute value of correlation coefficients, color corresponds to chi-square residuals. D. UMAP plots of re-grouped T cell subsets (n=20,930 cells), including naïve, CD8 effector, memory, effector memory, Treg, resting, and others. E. Correlation plots of over- or under-enrichment of each T cell subset (naïve, CD8 effector, memory, effector memory, Treg, resting, and others) from the WBCs of cancer patients with CTC positive, CTC low, and CTC negative, and healthy controls. Dot size corresponds to the absolute value of correlation coefficients, color corresponds to chi-square residuals. F. DPT cells highlighted on T cell subset UMAP plots, split by CTC status G. UMAP plot of mouse T cells, including single positive and DPT cells collected from Balb/c mouse splenocytes. H. Volcano plot depicting most differentially expressed genes in DPT cells versus all other T cells in mouse splenic T cells from G. I. Venn diagram showing the overlap between adhesion molecule genes expressed in total human T, mouse T, human DPT, and mouse DPT cells, and a list of 32 genes shared among 4 groups. J. Violin plots of mouse *Itgb1* mRNA expression in mouse T cells (CD4, CD8, and DPT) isolated from splenocytes, as measured by scRNA-seq. One-way ANOVA non-parametric (Kruskal-Wallis) test P values provided. K. Bar graphs of Cd29 (encoded by *Itgb1*) and Cd49d (encoded by *Itga4*) expression (mean fluorescence intensity, MFI) in mouse DPT, CD4, and CD8 cells isolated from Balb/c mouse splenocytes. One-way ANOVA P values provided. N=6 mice.

To further explore this phenomenon, we analyze specific subtypes of T cells. Here we identified 8 subsets of T cells based on their transcript expression, such as naïve T cells (*TCF7)*, CD8 effector (*NKG7*), memory and effector Memory (*IL7R*), and regulatory T cells (*FOXP3*) (**Figure 4D, Figure S6D**). While healthy controls had enriched effector memory T cells, CTC-positive cancer patients had over-represented naïve T cells (**Figure 4E, Figure S6E-S6F**). In contrast, CTC-low and -negative patients possessed a relatively high abundance of circulating effector and memory T cells (**Figure 4E**), which might be able to reject CTCs upon encountering them. Consistent with the literature ^42^, Tregs were slightly over-represented in cancer patients in our dataset analyses (**Figure 4E**).

Based on transcript expression, we identified DPT cells (∼500 cells) that were distributed into CD8 effector, memory (mostly CD4^+^), and naïve T subsets with a clear enrichment in CTC-positive patients (**Figure 4F, Figure S6G**). Using a public scRNA-seq database of peripheral blood cells from patients with triple-negative breast cancer ^43^, we extracted additional DPT cells for combined profiling with ours (a total of 1,454 cells), which also partially resemble the CD4 and CD8 T cells plotted in similar numbers (**Figure S6H**). Furthermore, we conducted scRNA-seq with sorted mouse splenic single-positive T and DPT cells, which form multiple subgroups with relatively distinct profiles from single positive T cells in the UMAP plot (**Figure 4G**). Compared to single-positive T cells, the mouse DPT cells show the strongest enrichment for immunosuppressive genes *Foxp3* and *Ctla4*, activation gene *Il2ra (Cd25)*, and integrin *Itgb1,* along with moderately enriched *Itga4* (**Figure 4H**).

In search of the molecular candidates contributing to interactive DPT-CTC clustering, we started with the KEGG cell adhesion and network genes (n=598) ^44^ and identified a list of 32 genes with overlapping expression among mouse and human T and DPT cells, including integrins *ITGB1* and *ITGA4*, *SELL, CD2, etc.* (**Figure 4I**). Notably, mouse *Itgb1* (Cd29) and *Itga4* (Cd49d), which together form integrin dimer VLA-4, were among the most enriched molecules in DPTs compared to single positive T cells (**Figure 4H-J, Figure S7A**). The higher expression levels of integrin proteins Cd29 and Cd49d were also observed in DPTs relative to single positive T cells (from mouse spleen and blood), as demonstrated via flow cytometry analyses (**Figure 4K, S7B-S7D**). Consistently, a similar but moderate enrichment of human *ITGB1* expression was also observed in blood DPTs versus CD4 and CD8 T cells, based on scRNA-seq data of breast cancer patients (**Figure S7E**).

To determine the functional importance of these candidate genes in heterotypic T cell interaction with tumor cells, we conducted genetic modulations by transient transfection of Jurkat T cells (or breast cancer cells) that stably express Cas9 with two individual synthetic guide RNAs (sgRNAs) for each gene. We first knocked out *ITGB1* and *SELL* in Jurkat cells and confirmed their depletion via flow cytometry (**Figure S7F**, left panels). As sgRNAs were unavailable for *CD2*, we alternatively knocked out *CD58*, the cognate ligand for CD2, in MDA-MB-231 breast cancer cells (**Figure S7F**, right panel).

We then performed a surrogate screening test for heterotypic T-tumor cell clustering *in vitro*, with 40,000 Jurkat cells (red) and 10,000 MDA-MB-231 cells (green) added to simulated cell suspensions in a poly-HEMA coated 96 well plate. As monitored by the IncuCyte time-lapse imaging of the 12-hour clustering, among all tested gene modulations, *ITGB1* knockout (KO) in Jurkat cells via two separate sgRNAs showed the strongest inhibition of heterotypic T-tumor cell interactions with differences shown as early as 3 hours (**Figure S7G-S7H**). One sgRNA for *SELL* KO in Jurkat cells also moderately reduced heterotypic interactions, whereas another *SELL* sgRNA mediated KO in Jurkat cells or *CD58* KO in MDA-MB-231 cells did not significantly impact heterotypic clustering (**Figure S7G-S7J**). Therefore, ITGB1 (VLA4 in partnership with ITGA4) became the top candidate to be further investigated in mouse models *in vivo*.

### VCAM1 is required for CTC-DPT clustering in a spontaneous metastasis model

To identify the ligand(s) in tumor cells that interact with the candidate receptors in DPT cells, we expanded the scRNA-seq data with periphery blood cells from breast cancer patients^45^ that include a subset of CTCs among the WBC populations (**Figure 5A**). From the heat map of predicted ligands in CTCs for the top adhesion molecules expressed in DPTs, we found two top adhesion molecules, *ICAM1* and *VCAM1* with higher expression in the CTCs from the patients with metastatic breast cancer than those from local disease (**Figure 5B-C**). Our previous work demonstrated that ICAM1 promotes cancer stemness and homotypic CTC cluster formation ^14^. Analyzing the mass spectrometry proteomic datasets of treatment-naive human breast tumors^46^ (N=122), we found a higher expression of VCAM1 protein in Black patients with breast cancer versus non-Black patients (**Figure 5D**). As VCAM1 is a known ligand of VLA-4 and is involved in breast cancer metastasis^47,48^, we hypothesized that VCAM1 contributes to tumor cell-T cell interactions.

**Figure 5.**
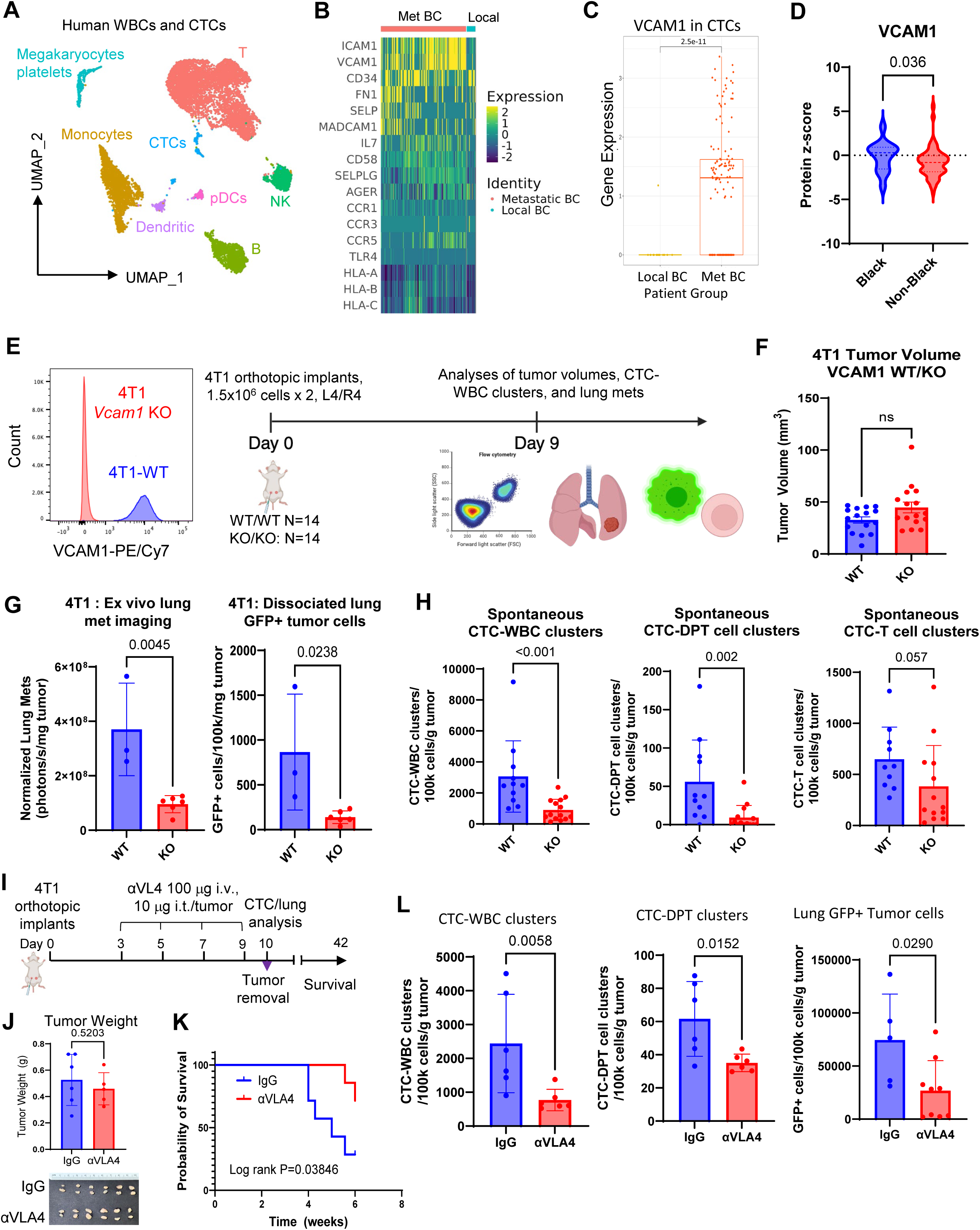
Targeting VCAM1-VL4 network inhibits spontaneous lung metastasis and CTC-DPT clusters *in vivo*. **A.** UMAP plot of CTCs and WBCs analyzed from a scRNA seq dataset of patients with breast cancer. **B-C.** Heatmap of ligand adhesion molecules (B) and bar graph of VCAM1 (C) in CTCs isolated from the patients with local and metastatic (Met) breast cancer (BC). **D.** Violin blot of proteomic VCAM1 in non-treated breast cancer tumors of Black and non-Black patients. Unpaired t-test, one-tailed, non-parametric. N = 14 for Black patients, 108 for Non-Black. **E.** Left panel: Flow histograms of mouse 4T1 tumor cells, WT and *Vcam1* KO. Right panel: Schematic depicting experimental design of orthotopic implants of eGFP+ 4T1 (WT and *Vcam1* KO) tumors in Balb/c mice and subsequent analyses of tumor burden, CTCs, and lung metastasis. **F.** Bar graphs of 4T1 primary tumor volumes, WT and *Vcam1* KO, on Day 9 (ns= not significantly changed, P>0.05) prior to eGFP immunogen triggered immune attacks in mice. N = 16 tumors. **G.** Bar graphs of lung metastatic signals of eGFP^+^ 4T1 cells detected via *ex vivo* fluorescence imaging of dissected lungs (left panel) and flow cytometry of dissociated eGFP+ cells from the mouse lungs (right panel). N = 3 for WT, 6 for KO. Unpaired, two tailed T test P values displayed. **H.** Bar graphs showing CTC-WBC clusters, CTC-T cell clusters, and CTC-DPT cell clusters in peripheral blood of mice with 4T1-NT control or VCAM1KO tumors. N = 11 for WT, 14 for KO. Unpaired, two-tailed t test P values displayed. **I.** Schematic of anti-VL4 neutralizing antibody (αVLA-4 Ab) treatment for orthotopic 4T1 breast tumors on days 3, 5, 7, 9. Blood/tissue harvests for primary tumors, CTCs, and lung metastasis analyses on Day 10 (N = 6). The primary tumor was removed via survival surgeries with the rest of mice (N=7 per group) for extended survival analyses until 6 weeks. **J.** Histogram graphs of 4T1 tumor weight and photos of dissected tumors of IgG control and αVLA-4 treated groups on Day 10. Unpaired two-tailed T test P = 0.197 (N=6) between two groups. **K.** Mouse survival after neoadjuvant treatment of IgG and αVLA-4 followed by a surgical removal of primary tumors on day 10 (N=7). Long rank P = 0.03846 for distinct survival of two groups of the mice by 6 weeks. Hazard ratio =4.435 (0.9842 to 20.75, 95% CI of ratio) when the IgG group is compared with αVLA-4 treated group. **L.** Bar graphs of blood-detected CTC-WBC clusters (left panel) and CTC-DPT clusters (middle manner), and the burden of lung tumor cells (right panel) detected and quantified via flow cytometry on Day 10, N = 6. Unpaired, two-tailed T test P values are shown.

To test the hypothesis, we employed the 4T1 mouse syngeneic model and patient CTC-derived CTC-205 model, both of which develop spontaneous metastasis to the lungs. We knocked out *Vcam1* in stably expressing Cas9 4T1 cells, implanted them into the mammary fat pads of Balb/c mice, and allowed tumors to grow with CTCs detected as early as 9 days to 16 days post-implantation before luciferase and/or fluorescent protein labels activated immune attacks shrink tumors (**Figure 5E)**. We harvested blood, lungs, and tumors, and assessed CTC-WBC clusters, immune infiltrates in the tumor and lungs, and metastatic burden in these animals.

While primary tumor burden was not significantly altered between the 4T1-WT and *Vcam1* KO groups (**Figure 5F**), significant decreases in metastatic signals were observed in the lungs of 4T1-*Vcam1*KO mice, as detected by biofluorescent imaging and flow cytometry (**Figure 5G**). Additionally, *Vcam1* KO in 4T1 cells reduced the formation of CTC-WBC clusters, nearly eliminating CTC-DPT cell clusters without altering other CTC-T cell clusters (**Figure 5H-J**). Interestingly, CTC clusters with granulocytic myeloid-derived suppressor cells (gMDSCs) also decreased in the blood of *Vcam1 KO-bearing* mice when other heterotypic clusters with B, monocytic MDSCs (mMDSCs) and neutrophils did not alter significantly (**Figure S8** and **S9A**).

In the primary tumors, *Vcam1* KO caused a significant decrease in infiltrated CD3^+^ T cells, coupled with increased gMDSCs (**Figure S9B**). A similar increase of gMDSCs was observed in the lungs of these KO tumor-bearing mice (**Figure S9C**), along with a decrease in circulating T cells, B cells, and mMDSCs in the periphery blood of these mice (**Figure S9D**). Together, these data show that Vcam1 on tumor cells is required for optimal CTC-DPT cell cluster formation and spontaneous lung metastasis in mice.

Next, we wanted to determine if the CTC-DPT interaction was dependent on TCR recognition of MHC molecules on CTCs, as genes involved in antigen recognition (*B2M, HLA* genes) were highly expressed in human and mouse DPT cells (**Figure 4H, I**). To address this question, we preincubated mouse splenic CD3 T cells with anti-MHC-I neutralizing antibodies and performed a clustering assay with 4T1 cells. Anti-MHC-I treatment did not interfere with the heterotypic cluster formation *in vitro* (**Figure S10A**).

To verify the role of VCAM1/VLA-4 interactions in human breast cancer, we profiled VCAM1 expression in cell lines and multiple patient-derived xenograft (PDXs) models (such as TN1 and CTC-derived model CTC-205) we generated ^33,38^. Our previous work identified VCAM1 was one of the top two enriched genes (along with ICAM1 and the stemness signature) in the lung metastases of PDXs compared to primary tumor cells ^14^. We further confirmed the higher expression VCAM1 in spontaneous metastases of PDXs (CTC-205) with minimal VCAM1 expression in primary tumors (**Figure S10B**).

Compared to the VCAM1^−^ tumor cells from the primary tumor xenograft (CTC-205), the VCAM1^+^ tumor cells sorted from the lung metastases (CTC-205-Met-VCAM1) promoted heterotypic clustering with wild-type (WT) Jurkat cells. However, this interaction was disrupted when the tumor cells were mixed with *ITGB1* KO Jurkat cells (**Figure S10C-S10D**). These findings suggest that the human tumor cell-T cell interactions also depend on a molecular network involving VCAM1 on the tumor cells and ITGB1 on T cells.

### Targeting the VLA-4 axis reduces CTC-DPT clusters and lung metastatic burden

To assess whether targeting VLA-4 could be translated into a therapy for treating metastatic breast cancer, we investigated the use of neutralizing antibodies. We hypothesized that anti-VLA-4 treatment would reduce CTC-DPT clusters and thus limit metastasis.

To test this, we implanted orthotopic 4T1 tumors into mice for neoadjuvant treatment with anti-α4-integrin (100 µg) via tail vein injection, plus 10 µg per tumor in the tumor bed, beginning on day 3 and continuing every other day for a total of 4 treatments before surgical resection of primary tumors on day 10 (**Figure 5I**). This regimen did not significantly affect the primary tumor size (**Figure 5J**). Remarkably, αVLA-4 treatment significantly extended the animal survival (**Figure 5K**). The blood and lung analyses on day 10 revealed decreased CTC-WBC clusters and CTC-DPT cell clusters, as well as the number of tumor cells in the lungs in the mice treated with αVLA-4 (**Figure 5L**). Notably, we observed no significant differences in the immune cell populations within circulation, the primary tumor, or the lungs between the control and treated groups (**Figure S11A-S11C**).

Collectively, these data suggest that blocking VLA4-mediated interactions and clustering of WBC/DPT and CTC offers a novel strategy to reduce the metastatic burden in breast cancer.

## Discussion

By integrating CTC analyses from a large cohort of breast cancer patients with mechanistic studies, our work uncovers a novel discovery of heterotypic CTC-WBC clusters containing a rare subset of DPT cells with unique features of exhaustion and immune suppression (TIM-3 and PD-1), thereby fostering tumor pluripotency and immune evasion. DPT cells are a rare subset of T cells and have been observed in association with many diseases, such as the periphery and tumor infiltrates of various cancers ^41,49–54^. We demonstrate that DPT-CTC clusters drive breast cancer metastasis via VLA4 (ITGA4/B1) - VCAM1 interactions. Notably, upregulated VCAM1 expression is observed in breast cancer of Black patients along with a higher frequency of heterotypic CTC-WBC clusters than white patients. That is positively associated with clinical outcome disparity of breast cancer, especially TNBC, with a disproportionately higher mortality risk for Black patients than for White and other patients ^55–59^, mainly due to metastasis ^60,61^.

To effectively prevent and eliminate metastasis, it is necessary to target primary tumors, CTCs (single, homotypic clusters, and diverse heterotypic clusters), and metastases in distant organs. In addition to targeting homotypic CTC clusters^6,14–21,62^, one of the pivotal therapeutic directions would be interfering with CTC-WBC clusters, especially the interaction with immune-suppressive DPT, through disruption of the VCAM1-VLA4 interactions and their related networks. Our studies demonstrate the proof of concept of targeting VLA-4 (α4) to prevent and eradicate CTC-WBC/DPT-mediated metastasis, ultimately improving breast cancer outcomes in Black and non-Black patients. Beyond cancer diseases, targeting VLA-4 has been safely and effectively extended to treat Crohn’s disease and multiple sclerosis^63^. Meanwhile, it is speculated that immune checkpoint inhibitors such as anti-PD-1 and anti-TIM3 would beneficially boost anti-tumor immunity for advantageous clearance of CTCs in close contact.

In addition to DPT cells, we have also detected other immune cells, such as neutrophils, monocytes, NK, and B cells in CTC clusters. The functional contribution and molecular mechanisms underlying these heterotypic CTC-WBC clusters can be investigated for therapeutic targeting. Neutrophils^64^ and monocytes are known to promote metastasis when they are incapable of removing tumor cells ^33,64–67^. Furthermore, NK cells^68^, platelets ^21,65,69,70^ and MDSCs ^33,71^ have also been shown to promote tumor cell seeding and metastasis. Additional strategies for specific targeting of CTCs and homotypic CTC-CTC clusters must be exploited to inactivate drivers of cancer stemness and CTC clustering, such as ICAM1^14^, via antibodies or antibody-drug conjugate, thereby blocking seeding and eliminating metastasis. Therefore, joint targeting efforts with cutting-edge technologies will accelerate the future reciprocal translation between the bench and the bedside.

## Materials and Methods

### Human sample analysis

Human blood sample collection and analyses were approved by the Northwestern University Institutional Review Board (IRB protocol # STU00203283 and STU00214936) following NIH guidelines for human subject studies. Written consent was obtained from all participants whose blood samples were analyzed for the study. CTCs (singles and clusters) of the blood samples collected in CellSave tubes were analyzed via CellSearch, and those in EDTA tubes were analyzed via flow cytometry, CellView, and ImageStream.

### Animal studies

All mice used in this study were housed in specific pathogen-free facilities, with regular diet, regular light/dark cycles, and regular ambient temperature and humidity in the Center for Comparative Medicine at Northwestern University. All animal procedures conformed to the NIH Guidelines for the Care and Use of Laboratory Animals (IACUC protocol IS00014098 and IS00021741) and were accepted by the Northwestern University Institutional Animal Care and Use Committees. For syngeneic models, female Balb/c mice aged 8-12 weeks (Jackson Laboratory) or female NSG mice aged 8-12 weeks (Jackson Laboratory) were randomized by age for orthotopic implantation with murine 4T1 cell line, human MDA-MB-231 cell line, or multiple triple negative breast cancer patient derived xenograft models.

### Sex as a biological variable

Our study mostly analyzed blood biospecimens from female patients, who account for over 99% of the patient population in breast cancer. Therefore, we exclusively examined female mice because the disease modeled is most relevant in females.

### Antibody treatments

Anti-VLA-4 (BioXCell, cat # BE0071) or IgG control antibody (BioXCell, cat # BE0090) were diluted to a final concentration of 500 µg/mL in PBS. Animals were injected with up to 100 µg of antibody via the tail vein and up to 10 µg of antibody into the tumor bed via subcutaneous injection. Each mouse received treatment every other day for 4 times.

### Breast tumor models and transfections

Murine 4T1 cells, human MDA-MB-231 and Jurkat cells were acquired from ATCC and regularly tested for mycoplasma using MycoAlert Mycoplasma Detection Kit (Lonza, cat # LT07-318). Patient derived xenograft model PDX-CTC-205 was generated from CTCs of a breast cancer patient, as described previously^38^. Early passages of cells (< 20 passages) were cultured in RPMI (4T1, Jurkat) or DMEM high glucose (MDA-MB-231) supplemented with 10% fetal bovine serum and 1% penicillin/streptomycin.

#### Cas9 transfection and CRISPR knockouts

For lentivirus production, HEK293T cells at 70-80% confluency were co-transfected with LentiCas9-EGFP (Addgene, plasmid #63592) and envelope vectors (pMD2.G and psPAX2) using the PEI reagent (Polysciences, cat # 23966). Supernatant was harvested 48 and 72 hrs after transfection, filtered with a 0.45 µm PES filter, centrifuged at 100,000 *g* for 2 hours, then aliquoted and stored in −80°C. 3 × 10^5^ MDA-MB-231, 5 × 10^5^ Jurkat, or 2 × 10^5^ 4T1 cells were cultured in 6 well plates and transduced with concentrated lentiviral supernatant (MOI of 1) using polybrene (8μg/mL). After 24 hours, the media containing lentivirus supernatant was replaced with fresh media. Cells were cultured 5 more days then eGFP+ cells were separated using cell fluorescent activated cel sorting.

To generate CRISPR/Cas9 mediated knockouts, TrueGuide synthetic gRNAs (ThermoFisher Scientific) were resuspended in 1X TE buffer at a concentration of 100 µM. Stock solution of gRNA was stored at −20°C until further use. The day before transfection with gRNA, Cas9 expressing cells were plated in a 6 well plate at a concentration of 250,000 cells/well. The next day, 37.5 pmol of gRNA was mixed with 125 µL of Opti-MEM I Reduced Serum Medium (ThermoFisher Scientific, cat # 31985062) per transfection well. 7.5 µL of Lipofectamine CRISPRMAX Cas9 Transfection Reagent (ThermoFisher Scientific, cat # CMAX00001) was added to 125 µL Opti-MEM I Medium per transfection well and incubated at room temperature for 5 minutes. The diluted transfection reagent solution was added to the tube containing the gRNA solution and mixed by pipetting. This solution was incubated for 15 minutes at room temperature. 250 µL of the gRNA/transfection reagent complex was added to each of the wells. Cells were allowed to grow for 2-3 days, upon which they were harvested, stained for the protein targeted by the gRNA, and analyzed by flow cytometry. Cells negative for the protein were sorted using a BD FACSAria sorter, replated, and propagated *in vitro*. Cells were then stained for flow cytometry and single cells were sorted into 96-well plates to generate clones. Knockout efficiency of clones was determined via protein analysis, and individual and pooled clones were sorted for downstream use.

### CTC analyses by CellSearch and flow cytometry

#### CellSearch

The CellSearch System (Menarini Silicon Biosystems) is semiautomated for blood sample processing, enrichment of EpCAM+ epithelial CTCs using the Epithelial Cell Kit, and subsequent four-channel immunofluorescence staining, as described by Cristofanilli et al. CTCs are specified by combining three routine channels of DAPI+, cytokeratin+, and CD45-. Single CTCs, homotypic CTC clusters, and heterotypic CTC clusters were manually verified and counted.

#### Flow cytometry

Patient blood samples were collected in EDTA Vacutainer tubes (Becton Dickinson) and stored on ice for processing within 24 hours after collection. Samples were spun at 300 x *g* for 10 minutes with no brake to remove plasma, which was stored at −80°C. Remaining blood cells were subject to 2-4 rounds of red blood cell lysis using Red Blood Cell Lysing Buffer Hybri-Max™ (Sigma-Aldrich, cat # R7757). White blood cells were washed in PBS and resuspended in PBS + 2% FBS. Cells were blocked using mouse IgG (Bio-Rad, cat # PMP01) for 10 minutes on ice and stained with antibodies for EpCAM, CD45, CD3, CD4, CD8, CD19, CD11b, CD14, CD16, CD56, and CD66b on ice, protected from light, for 30 minutes. Cells were washed with PBS and resuspended in PBS + 2% FBS, stained with DAPI or Live/Dead Fixable Blue Dye (ThermoFisher Scientific, cat # L23105), and analyzed on a BD Fortessa analyzer. For ImageStream analysis, samples were run on a Cytek Amnis ImageStream^x^ Mk II imaging flow cytometry (Cytek Biosciences) and analyzed using the IDEAS image analysis software version 6.2.

### Splenocyte and T/DPT cell collection

Female Balb/c mice were sacrificed by CO_2_ asphyxiation followed by cervical dislocation. Spleens were removed and forced through a 70 µm mesh filter (Fisher Scientific, cat # 22363548) using the plunger of 3 mL syringe (Fisher Scientific, cat # 14823435). Cells were washed in PBS, spun at 300 *g* for 5 minutes and resuspended in red blood cell lysis buffer. Cells were incubated in lysis buffer on ice for 5 minutes, washed in PBS to stop the reaction, and spun at 300 *g* for 5 minutes. Cells were resuspended in PBS + 2% FBS + 20 mM EDTA, filtered through a 40 µm mesh filter (Fisher Scientific, cat # 22363547), and counted. To isolate T/DPT cells, the MojoSort Mouse CD3 T Cell Isolation Kit (Biolegend, cat # 480024) was used. Briefly, unlysed splenocytes were incubated with the Biotin-Antibody Cocktail for 10 minutes on ice, followed by the Streptavidin Nanobeads for 10 minutes on ice. Cells of interest were separated using the MojoSort Magnet (Biolegend, cat # 480019) for 2 separations. Cells were washed in PBS and resuspended in PBS + 2% FBS + 20 mM EDTA. To further isolate specific T cell populations (CD4, CD8, or DPT), cells were blocked with TruStain FcX (anti-mouse CD16/32) Antibody (Biolegend, cat # 101319) for 10 minutes on ice and stained with antibodies against mouse CD45, CD3, CD4 and CD8. Samples were sorted on a BD FACSAria SORP system or a Miltenyi MACSQUANT Tyto Cell Sorter using MACSQUANT Tyto Cartridges HS (Miltenyi Biotec, cat # 130121549) for downstream use.

### Clustering Assay

For cell lines, 96-well plates were coated with poly-HEMA (Sigma Aldrich, cat # P3932) overnight at room temperature, then washed with PBS. For PDX cells, 96-well plates were coated in collagen I, bovine (Corning, cat # 354321), diluted 1:100 in water, for 1 hour at room temperature, then washed with PBS. Cells were stained with PKH67 Green Fluorescent Cell Linker (Sigma Aldrich, cat # PKH67GL) or Incucyte Cytolight Red (Sartorius, cat # 4706) according to manufacturer’s instructions. Single cell suspensions of 4T1, MDA-MB-231, PDX cells, or immune cells were added to the wells at a final volume of 200 µL. Plates were imaged using the Incucyte ZOOM Live Cell Imaging System (Essen Biosciences) and analyzed using the Incucyte ZOOM Software. For the metastasis lung colonization assay, after clustering, cells were gently collected into EasyTouch U-100 insulin syringe (MHC Medical Products, cat # 829155) so as not to disturb clusters, and injected into the tail veins of Balb/c mice.

### Bioluminescent Imaging

Animals were injected i.p. with 100 µL D-luciferin at 30 mg/mL (Gold Biotechnology, cat # 115144359) and anesthetized using isoflurane, and then placed into the In Vivo Imaging System Lago X (Spectral Instruments Imaging) and bioluminescence was detected with an acquisition time up to 5 minutes. Images were analyzed using the Aura In Vivo Imaging Software (Spectral Instruments Imaging). Signal is reported as normalized total flux (photons/second).

### Immunohistochemistry

Tissues were fixed in 4% paraformaldehyde for 24-48 hours, and then transferred to 70% ethanol solution. Tissue was embedded in paraffin blocks, cut into 4 µm thick sections, and processed for hematoxylin & eosin or IHC staining by the Mouse Histology and Phenotyping Laboratory at Northwestern University.

### Cell isolation and single-cell RNA sequencing library preparation

Bulk white blood cells or mouse splenic T cells were processed as described above and CD45+ single cells or mouse T/DPT cells were sorted into tubes of PBS + 2% FBS on a BD FACSAria sorter. For 4T1 clustering samples, 4T1-DPT or 4T1-splenocyte clusters generated as described above and collected, washed with PBS, and clusters were gently disturbed to generate single cell suspensions before proceeding to next step. For experiments with multiplexing, sorted cells were blocked with Human TruStain FcX (Fc Receptor Blocking Solution (BioLegend, cat # 422301) or TruStain FcX (anti-mouse CD16/32) Antibody (Biolegend, cat # 101319) for 10 minutes on ice, followed by staining with TotalSeq-C anti-human or anti-mouse Hashtag Antibodies for 30 minutes on ice. Cells were washed 3 times in Cell Staining Buffer (BioLegend, cat # 420201) and resuspended in PBS + 2% FBS in a single cell suspension. The concentration and viability of the single cell suspensions were measured using a Nexcelom Cellometer Auto 2000 with ViaStain AOPI staining solution (Nexcelom, cat # CS2-0106). Cells were loaded onto a 10X Chromium Controller for GEM generation followed by single cell library construction using 10X Chromium Next GEM Single Cell 5’ Library and gel bead kit v1.1 (10X Genomics, cat # PN-1000165) according to the manufacturer’s protocol. Quality control of the libraries was performed using an Invitrogen Qubit DNA high sensitivity kit (ThermoFisher Scientific, cat # Q32851) and Agilent Bioanalyzer high sensitivity DNA kit (Agilent, cat # 5067-4626). The libraries were pooled in equal molar ratio and sequenced on an Illumina HiSeq 4000 using sequencing parameters indicated by the manufacturer (Read 1: 26 cycles, i7 index: 8 cycles, Read 2: 91 cycles).

### Generation of single-cell gene expression matrices

Demultiplexing, alignment, and gene quantification were all performed using Cell Ranger (10X Genomics, version 7.0.1). The run data was demultiplexed using cellranger mkfastq and the resulting FASTQ files were aligned and counted using the 10X Genomics pre-built human (refdata-gex-GRCh38-2020-A) or mouse (refdata-gex-mm10-2020-A) reference genomes.

### Quality control, cell-type clustering, and major cell type identification

For each sequencing run, 5% quantiles of genes expressed and unique molecular identifiers were calculated. Cells expressing less than 500 genes and UMIs and greater than the 90^th^ percentile were excluded from analysis. Furthermore, cells with greater than 10% mitochondrial content were also excluded. Datasets were integrated to control for batch effects among multiple samples and sequencing runs. Dimensionality reduction and unsupervised cell clustering was performed using methods from the Seurat software suite (version 4.3.2). The results are presented using Uniform Manifold Approximation and Projection (UMAP) and 19 distinct clusters were originally identified. After gene expression analysis, Seurat-determined clusters were collapsed into biologically relevant clusters that align with cell type. Cell types were identified by expression of CD3, CD14, CD16, CD19, NCR1, and FCGR2A. T cell clusters were split from the main dataset and analyzed separately. To identify DPT cells, cells that had normalized RNA expression of both CD4 and CD8A from this dataset and from a public dataset (GSE169246) were selected. Additionally, 1000 random CD4^+^CD8A^−^ cells and CD4^−^CD8A^+^ from each dataset were selected to generate a T/DPT only dataset for analysis.

### Statistical Analysis

GraphPad Prism 10 was used to calculate statistical tests. For all tests, p < 0.05 was considered significant. R (version 4.4.0) and packages Seurat (4.4.2), dplyr (1.1.4), tidyverse (2.0.0), ggplot2 (3.5.1), survminer (0.4.9), enrichR (3.3), and survival (3.7-0) were used for data analysis and to generate figures. FlowJo software (version 10.10.0) was used to generate flow cytometry plots and for data analysis. All graphs represent mean ± standard deviation unless otherwise specified.

## Data Availability

scRNA-seq datasets are available from the corresponding author upon reasonable request. Datasets GSE169246 and GSE139495 are available at: https://www.ncbi.nlm.nih.gov/geo/query/acc.cgi?acc=GSE169246 and https://www.ncbi.nlm.nih.gov/geo/query/acc.cgi?acc=GSE139495. Human spleen immunofluorescence images are based upon data generated by the HuBMAP Program^37^: https://hubmapconsortium.org.

## Acknowledgements

This work has been supported by the core facilities at Northwestern University including the Robert H. Lurie Comprehensive Cancer Center CTC Core, the Flow Cytometry Core, the Pathology Core, the Mouse Histology and Phenotyping Laboratory, and the Center for Comparative Medicine. We are deeply grateful to the clinical team and the patients who consented to donate their biospecimens. We thank all team members in the Liu Cluster and collaborative laboratories of Drs. Deyu Fang, Yan Liu, Jae Choi at Northwestern University and Dr. Clive H Wasserfall laboratory at the University of Florida. This project has been partially funded by Department of Defense grant W81XWH-20-1-0679 (H. Liu); NIH/NCI grants R01CA245699, R01AI167272, UG3CA256967, R01CA298232 (H. Liu) and Fellowship T32 CA009560 (D. Scholten); Chan Zuckerberg Biohub Chicago (H. Liu); American Cancer Society CSCC-Team-23-980420-01-CSCC (H. Liu); the Lurie Cancer Center IDH (H. Liu); Lynn Sage Breast Cancer Research Foundation (H. Liu, N. Dashzeveg, and L. El-Shennawy); and Northwestern University Pharmacology start-up grant (H. Liu).

## Author Contributions

D.S. and H.L. conceived the idea presented here and wrote the manuscript. D.S. completed the experiments and data analyses with support and help by L.E., Y.J., Y. Z., E.H., C.R., A.D.H, H.F.A, F.T., and N.D. Senior authors H.L., D.F., and M.C., C.H.W., W.G., J.L. supervised experimental planning and implementation. L.E. assisted with flow cytometry analyses of immune cells (WBCs) and CTCs (singles and clusters). L.E, Y.J., H.F.A., and N.D. helped the animal work *in vivo*. Y.Z. and C.Z collected CTC data from patients. E.H assisted with blood sample processing and data analyses. A.D.H and F.T., and Y.S facilitated the analysis of single-cell and proteomic data.

## Inclusion and Diversity

We are committed to fostering an inclusive research environment. Our study design and data analyses were structured to minimize biases and ensure equitable representation across relevant participant groups. Where possible, we stratified results to address biological and clinical variables such as sex, race (Asian, Black, White, and others), and ethnicity, respecting privacy and ethical constraints. Researchers from diverse backgrounds and expertise were involved throughout the project, reflecting our commitment to multidisciplinary collaboration and inclusive scientific practices.

## Conflict of interest statement

Huiping Liu, Deyu Fang, and Andrew D. Hoffmann are scientific co-founders and equity shareholders of Exomira Medicine Inc, whose business is not currently related to the content of this manuscript. David Scholten and Huiping Liu have a pending patent application relevant to the manuscript to be submitted. Massimo Cristofanilli has the following financial relationships to disclose: (1) serving as consultant for AZ, Celcuity, Menarini-Stemline, Repare Therapeutics, Olaris, Syantra, BriaCell, Datar Cancer Genomics, Biotheryx; (2) received grant or research support from AZ, Celcuity; (3) received honoraria from AZ, Merck, Iylon, Menarini-Silicon Biosystem, Datar Cancer Genomics; and (4) participated in Speaker Bureau of Pfizer.

