## Supplemental Figures for "Rare Subset of T Cells Form Heterotypic Clusters with Circulating Tumor Cells to Foster Cancer Metastasis"

Supplementary Figures S1-11

### Figure S1

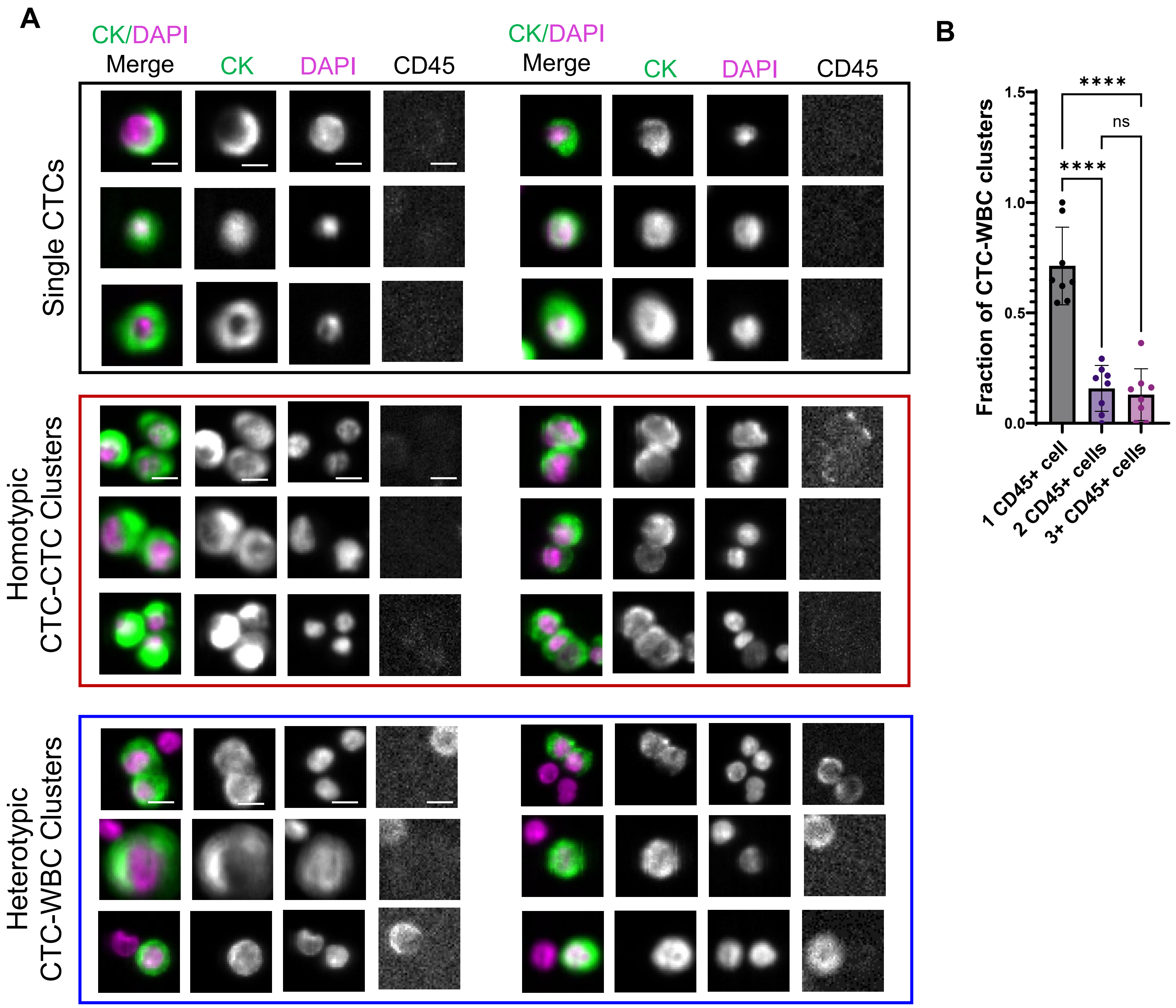


**Figure S1. Human CTCs (singles and clusters) from breast cancer patients detected via CellSearch.**

**A.** Representative images of CTCs, singles, homotypic CTC- CTC clusters, and heterotypic CTC-WBC clusters with merged or single channels of cytokeratin (CK, green), DAPI (magenta), and CD45. Top panels: Single CTCs. Middle panels**.** Homotypic CTC clusters. Bottom panels: Heterotypic CTC-WBC clusters.

**B.** Frequency of immune cell or WBC counts per CTC-WBC cluster. One-way ANOVA with Tukey's multiple comparison test; ****, $p < 0.001$. N = 8 CellSearch scans (patient biospecimens).

### Figure S2

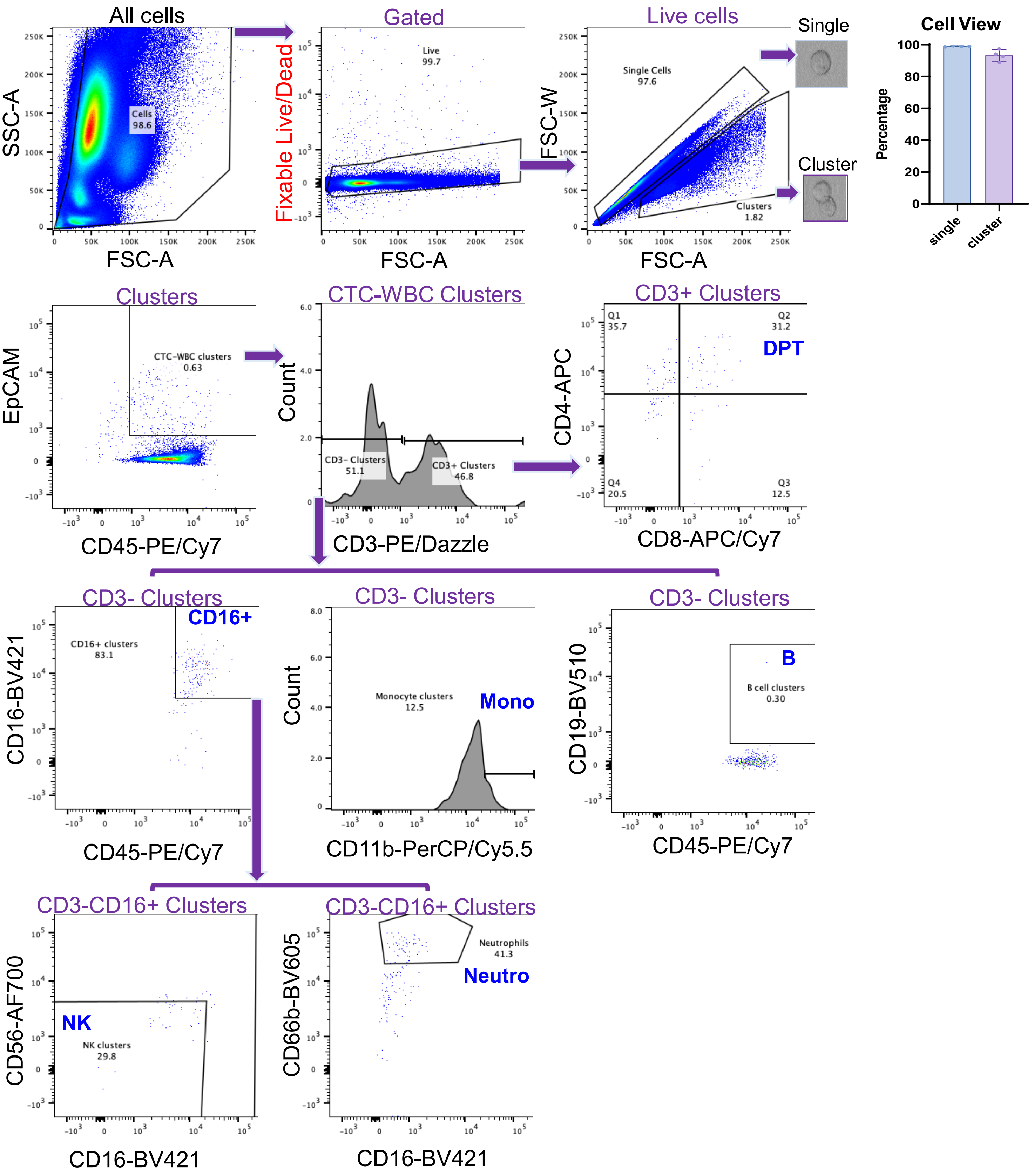


**Figure S2. Representative gating of CTC-WBC clusters.** Single cells and clusters were gated based on forward/side scatter to distinguish their distinct sizes at an accuracy of 99% and 95%, respectively, as validated by BD CellView. WBC-CTC clusters were gated on CD45+EpCAM+, with $CD3^{+}$ T cell clusters, other $CD3^{-}$ clusters for B cells ($CD19^{+}$); NK cells ($CD16^{+}CD56^{+}$); neutrophils ($CD16^{+}CD66b^{+}$); and monocytes ($CD11b^{+}CD16^{+/-}CD14^{+/-}$).

### Figure S3

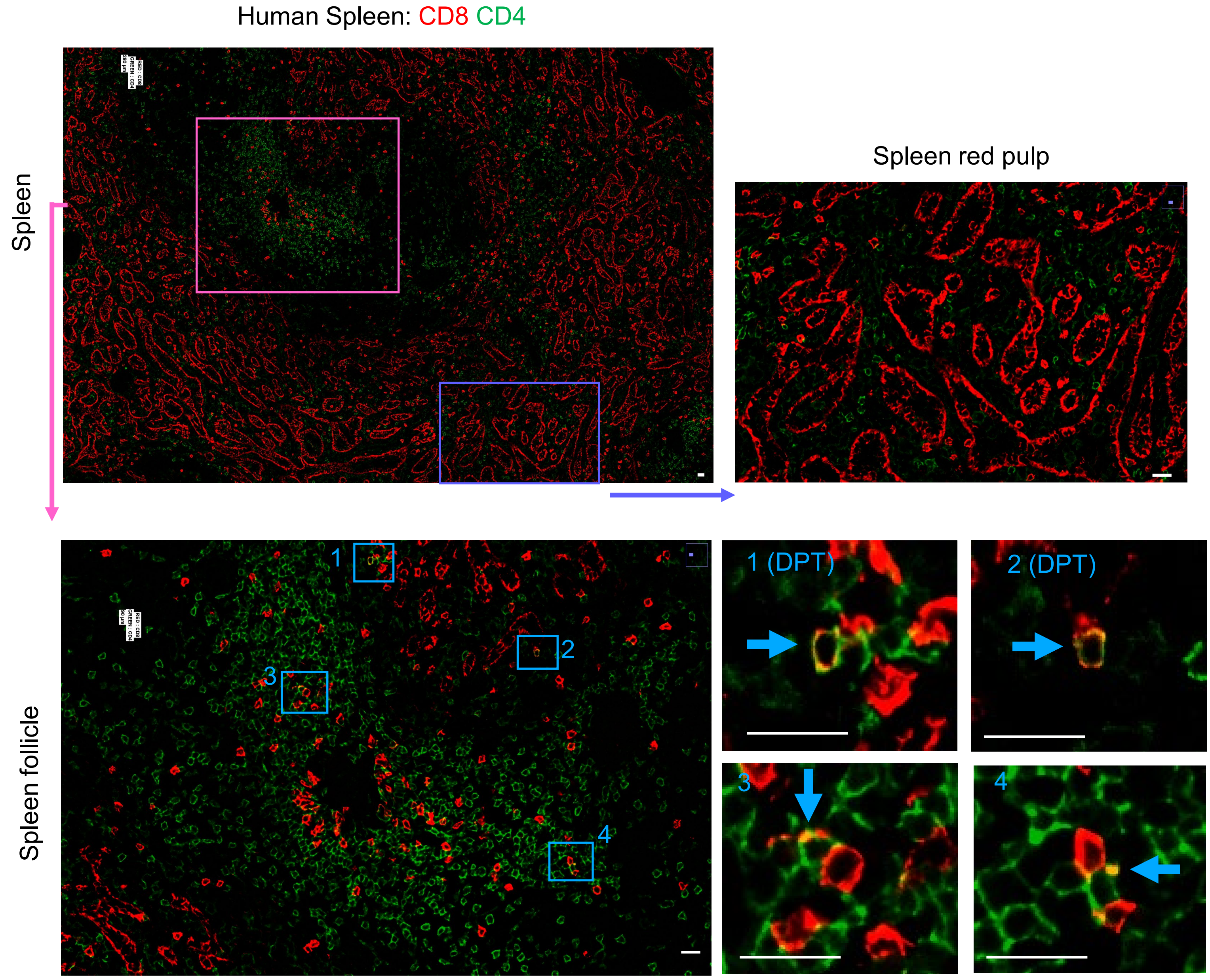


**Figure S3. CD4 and CD8 immunofluorescence staining of human spleens with DPTs**.
**Top left panel**: a low magnification image of the human spleen with chosen regions of follicle (pink borders) and red pulp (purple borders) stained with CD4 (green) and CD8 (red) antibodies.
**Top right panel:** zoom-in image of the red pulp insert with no detected DPTs.
**Bottom panels:** zoom-in image of the follicle insert with 4 highlighted inserts (blue) showing yellow surface of double positive staining of 2 DPTs in inserts 1 and 2 (blue arrows pointed).
Scale bars = 50 μm.

### Figure S4

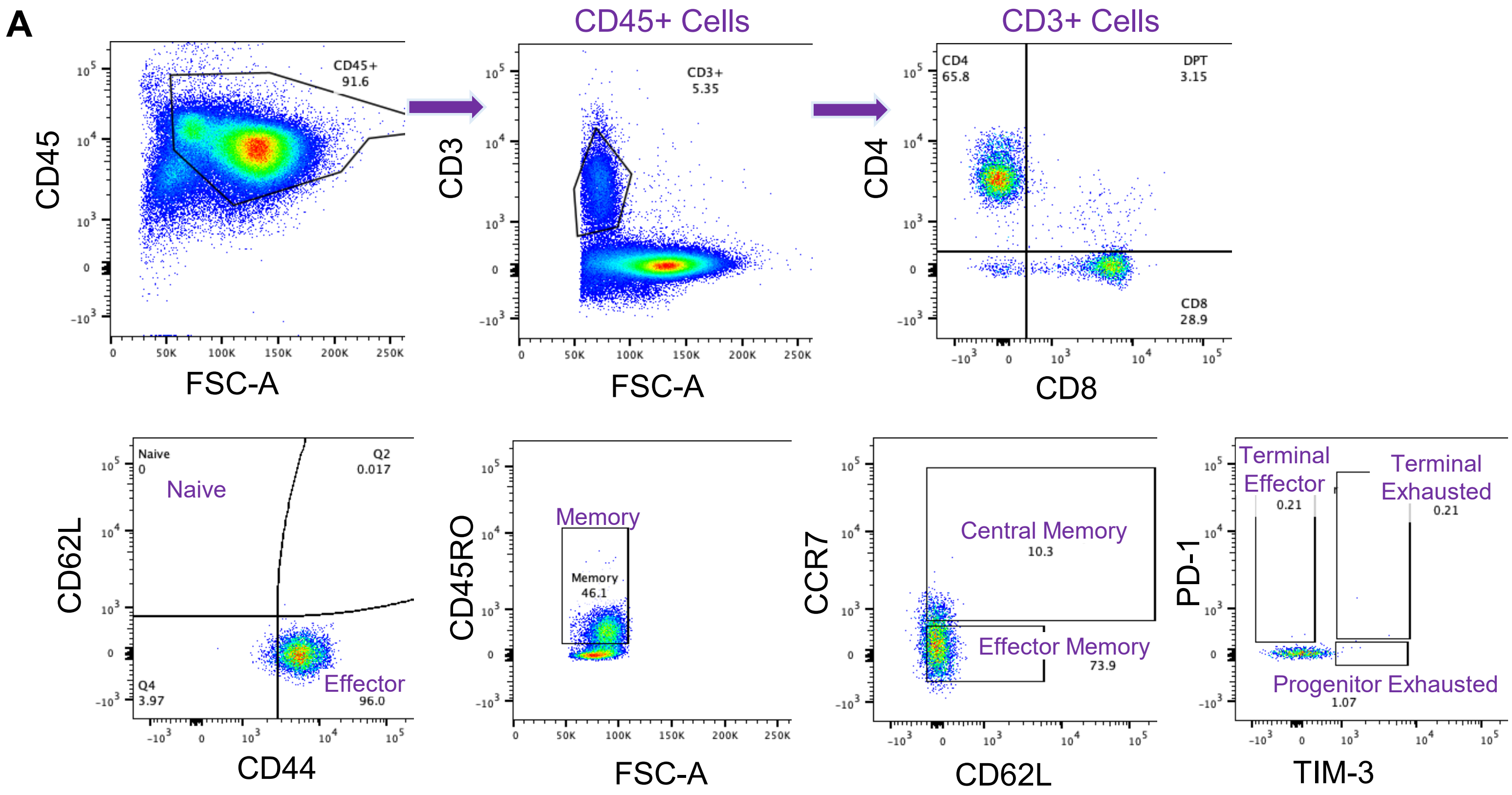


**Figure S4. Phenotypic characterization of human circulating DPT cells versus CD4⁺ and CD8⁺ single positive T cells in WBCs of patients with breast cancer.**

**A.** Representative gating of T/DPT cell subsets, including naïve ($CD62L^{+}CD44^{-}$), memory ($CD45RO^{+}$), central memory ($CD45RO^{+}CCR7^{+}$), effector memory($CD45RO^{+}CCR7^{-}$), terminal effector ($PD\text{-}1^{+}TIM3^{-}$), progenitor exhausted ($TIM3^{+}PD\text{-}1^{-}$), terminal exhausted cells ($PD\text{-}1^{+}TIM3^{+}$).

### Figure S5

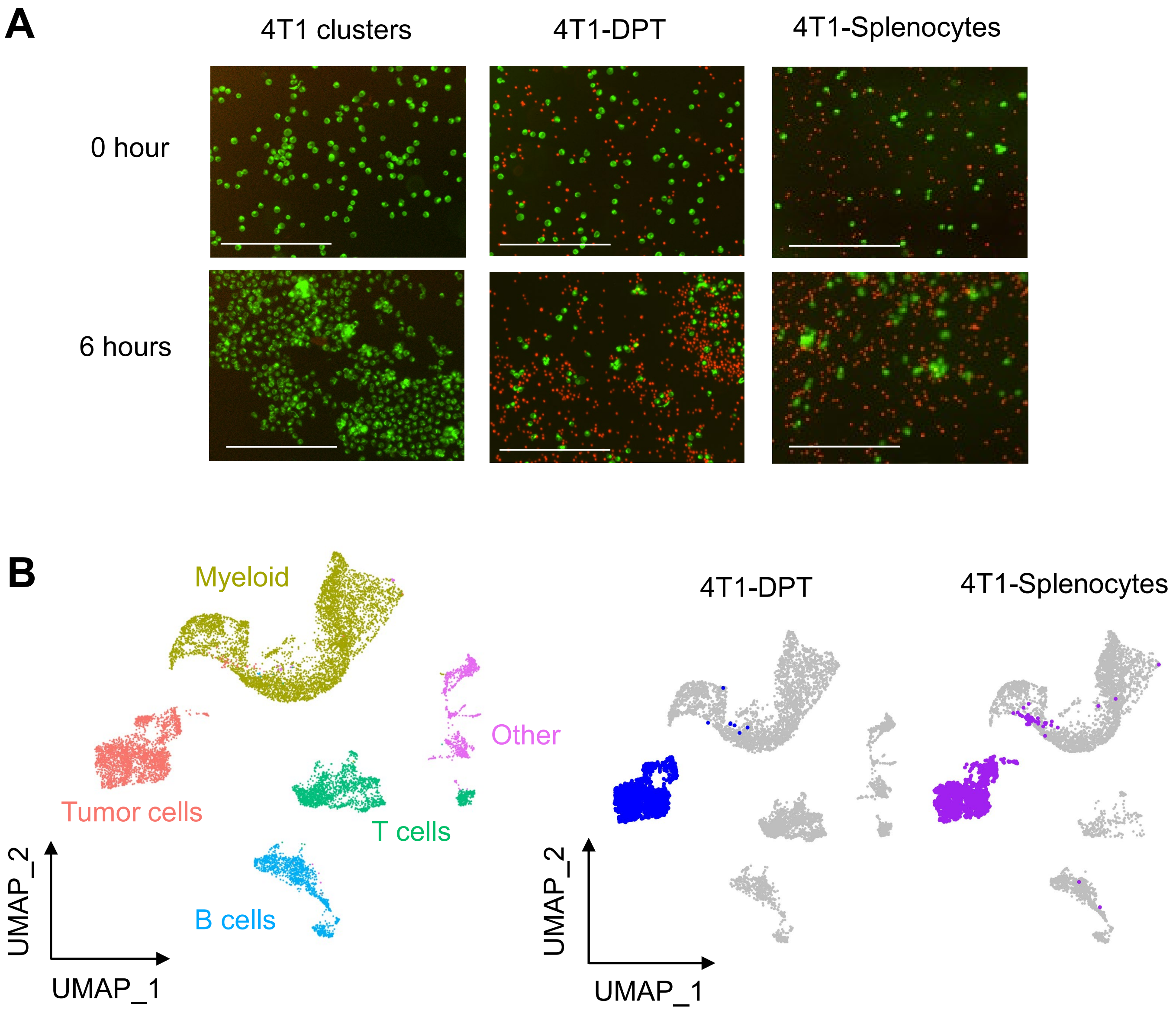


**Figure S5. 4T1-DPT and 4T1-splenocyte interactions influence tumor cells.**

**A.** Representative images of 4T1 tumor cell only-aggregated homotypic clusters (left panels), 4T1-DPT heterotypic interactions and clusters with DPT cells sorted from tumor-bearing splenocytes (middle panels), and 4T1- splenocytes (unsorted control, right panels) after 6 hours of clustering at 37°C. Green, 4T1 cells labelled by PKH67. Red, DPT cells and splenocytes labeled by Cytolight Red.

**B.** UMAP of 4T1-DPT cells and 4T1-splenocytes collected after 6 hours incubation (left). UMAP of tumor cells colored by coincubation cell type (right), blue = DPTs and magenta = splenocytes for co-incubation.

### Figure S6

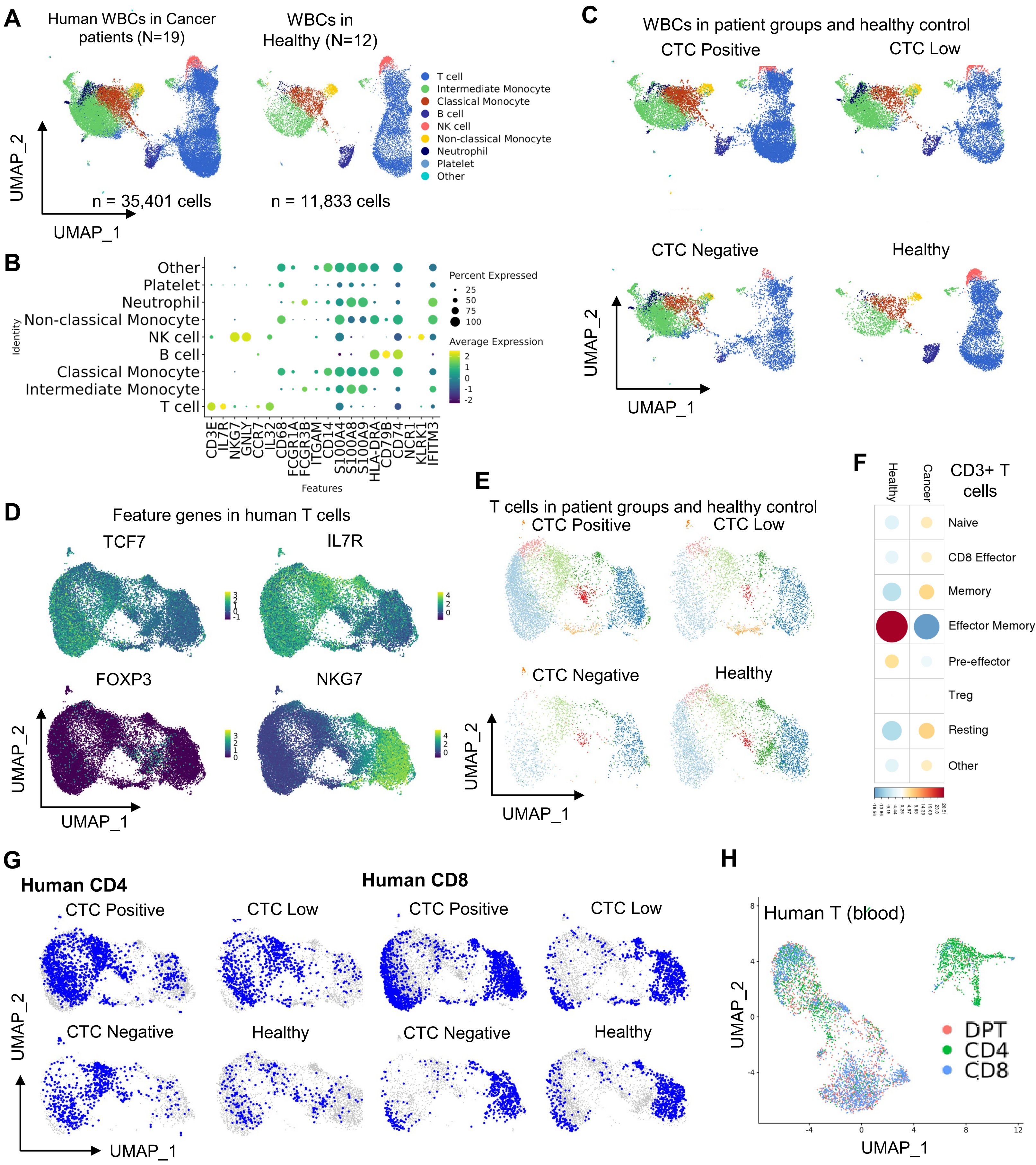
A
Human WBCs in Cancer patients (N=19)
WBCs in Healthy (N=12)
T cell
Intermediate Monocyte
Classical Monocyte
B cell
NK cell
Non-classical Monocyte
Neutrophil
Platelet
Other
UMAP_2
UMAP_1
n = 35,401 cells
n = 11,833 cells
B
Other
Platelet
Neutrophil
Non-classical Monocyte
NK cell
B cell
Classical Monocyte
Intermediate Monocyte
T cell
Identity
Percent Expressed
25
50
75
100
Average Expression
CD3E
IL7R
NKG7
GNLY
CCR7
IL32
CD68
FCGR1A
FCGR3B
ITGAM
CD14
S100A4
S100A8
S100A9
HLA-DRA
CD79B
CD74
NCR1
KLRK1
IFITM3
Features
C
WBCs in patient groups and healthy control
CTC Positive
CTC Low
CTC Negative
Healthy
D
Feature genes in human T cells
TCF7
IL7R
FOXP3
NKG7
E
T cells in patient groups and healthy control
CTC Positive
CTC Low
CTC Negative
Healthy
F
Healthy
Cancer
CD3+ T cells
Naive
CD8 Effector
Memory
Effector Memory
Pre-effector
Treg
Resting
Other
G
Human CD4
Human CD8
CTC Positive
CTC Low
CTC Negative
Healthy
H
Human T (blood)
DPT
CD4
CD8

**Fgure S6. scRNA-seq analysis of human WBCs and DPT cells**

**A.** UMAP plots of WBCs collected from breast cancer patients (left panel, N = 19 patients, with 35,401 cells) and healthy controls (right panel, N =12, with n = 11,833 cells). Cell populations are indicated in distinct colors.

**B.** Dot plot of representative markers used to annotate broad immune cell subsets in human WBCs. Dot size corresponds to the percent of expressing cells, and color corresponds to the expression levels.

**C.** UMAP plots of WBCs split by CTC status, including breast cancer patients with CTC positive (≥5 CTCs), CTC low (1-4 CTCs), and CTC negative (0 CTCs) per 7.5 mL blood, and healthy controls (non-cancer). Dot size corresponds to the absolute value of correlation coefficients, and color corresponds to chi-square residuals.

**D.** Feature plots depicting representative genes used to annotate T cell subsets, including *TCF7*, IL7R, FOXP3, and NKG7.

**E.** Human T cell subset UMAP plots of patient WBCs split by CTC status, including CTC positive, CTC low, and CTC negative, and healthy controls.

**F.** Correlation plots depicting chi-squared test residuals to determine over- or under-enrichment of each T cell subset from the T cells of cancer patients vs. healthy controls. Dot size corresponds to the absolute value of correlation coefficients, and color corresponds to chi-square residuals

**G.** CD4 (left) and CD8 (right) T cell distribution in the UMAP plots within four groups as split by CTC status.

**H.** UMAP plot of combined human circulating T (CD4 and CD8) and DPT cells from our blood cell dataset and a public dataset (*Zhang et al.*), which show partially overlapping profiles between DPT and single positive-cell controls.

### Figure S7

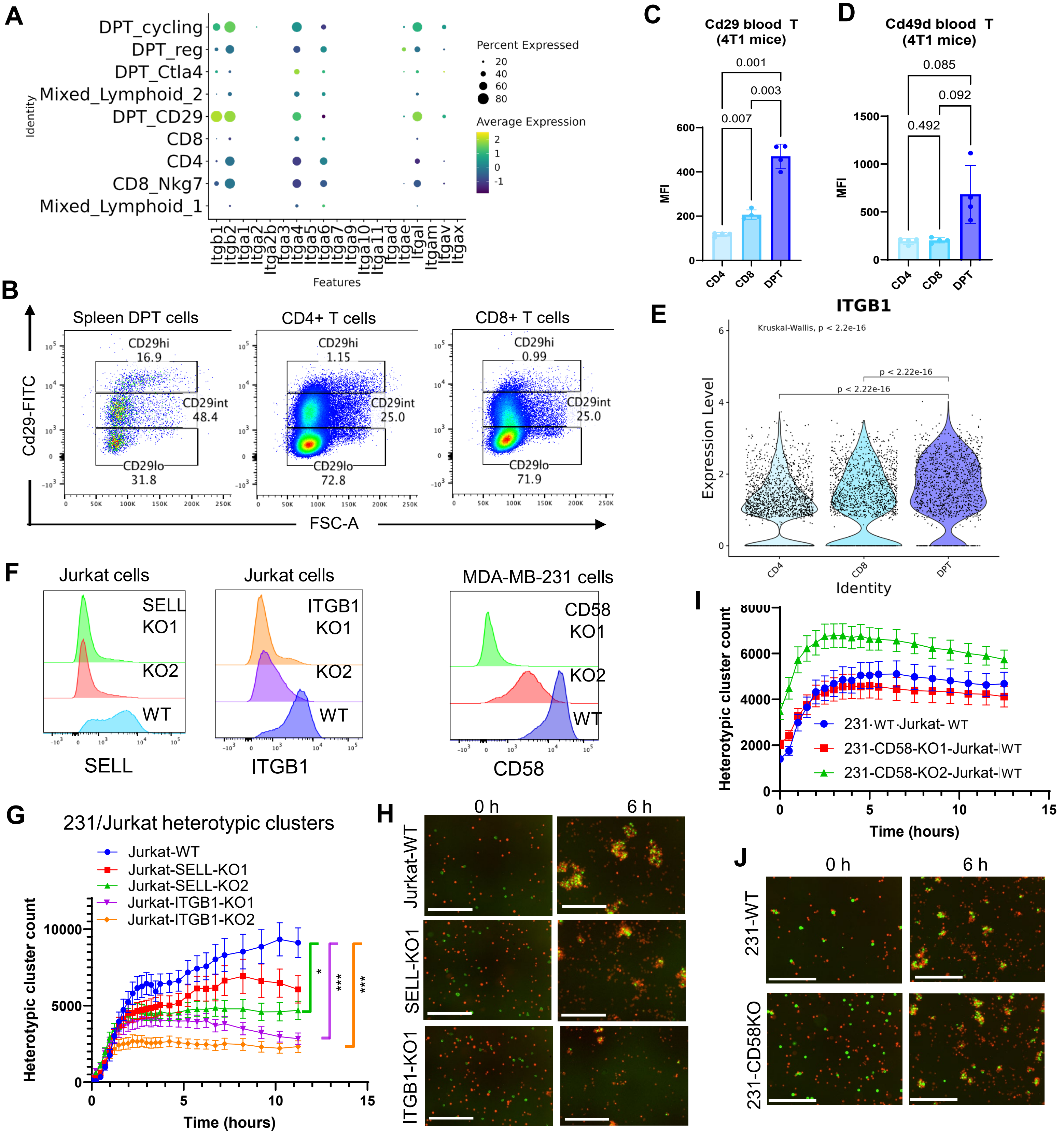


**Figure S7. Integrins and VCAM1 as mediators of T cell-tumor cell clustering. A.** Circle plot of integrin alpha and beta gene expression (mRNAs) in mouse splenic T and DPT cells. **B.** Flow plots of Cd29 protein expression in DPT, CD4, and CD8 cells. **C-D.** Bar graphs of Cd29 (C) and Cd49d (D) expression (mean fluorescence intensity, MFI) in mouse DPT, CD4, and CD8 cells from WBCs. One-way ANOVA P values (N=6 mice). **E.** Violin plots of human ITGB1/CD29 expression in DPT, CD4, and CD8 T cells. **F.** Flow histograms of Jurkat-Cas9-KO cells (SELL and ITGB1) and MDA-MB-231-Cas9-KO cells (CD58). G. Curves of heterotypic cluster counts of Jurkat-Cas9 cells, control (NT) or genetic knockout of *SELL* or *ITGB1,* and MDA-MB-231 tumor cells. Graphs represent mean ± standard deviation. Two-sided unpaired t-test. *, $p < 0.05$; ***, $p < 0.005$. N = 6 biological replicates. **H.** Representative images of G. **I.** Curves depicting heterotypic interactions between WT Jurkat cells and MDA-MB-231-CD58KO cells. Graphs represent mean ± standard error of the mean. N = 6 biological replicates. **J.** Representative images of the curves from I. Green, MDA-MB-231 cells (231); red, Jurkat cells. Scale bars = 300 µm.

### Figure S8

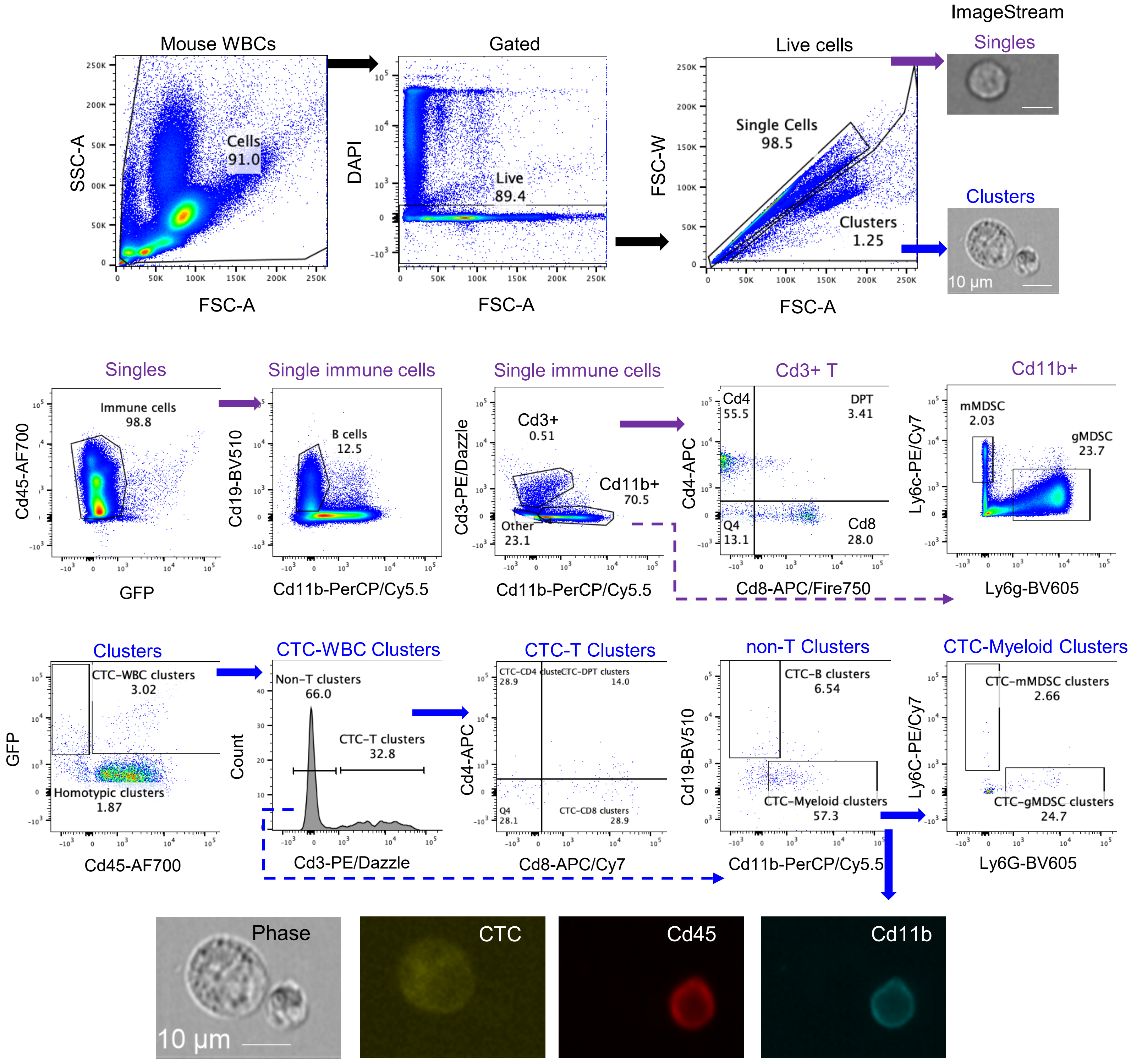


**Figure S8. Gating strategies for identifying mouse white blood cell (WBC) lineages in singles and CTC-WBC heterotypic clusters.** The sequential gates include the top row: mouse WBCs, live cells (DAPI-), singles and clusters. Second row: single cells, single immune cells ($CD45^+ GFP^-$), $Cd3^+$ T cells, $Cd4^+$ and $Cd8^+$ single positive T cells and DPT cells; $Cd19^+$ B cells, Cd $11b^+$ myeloid cells, mMDSC ($Ly6c^+$), and gMDSC ($Ly6g^+$). The third row: clusters, CTC-WBC clusters (eGFP+Cd45+), CTC-T clusters (Cd3+), non-T clusters (Cd3-), CTC-B clusters (CD19+Cd11b-), CTC-myeloid clusters (CD19-CD11b+), CTC-mMDSC clusters (Ly6c+Ly6G-), and CTC-gMDSC clusters (Ly6C-Ly6G+). The bottom row: ImageStream images of CTC-WBC cluster with a CTC and a CD45+CD11b+ WBC in the cluster.

### Figure S9

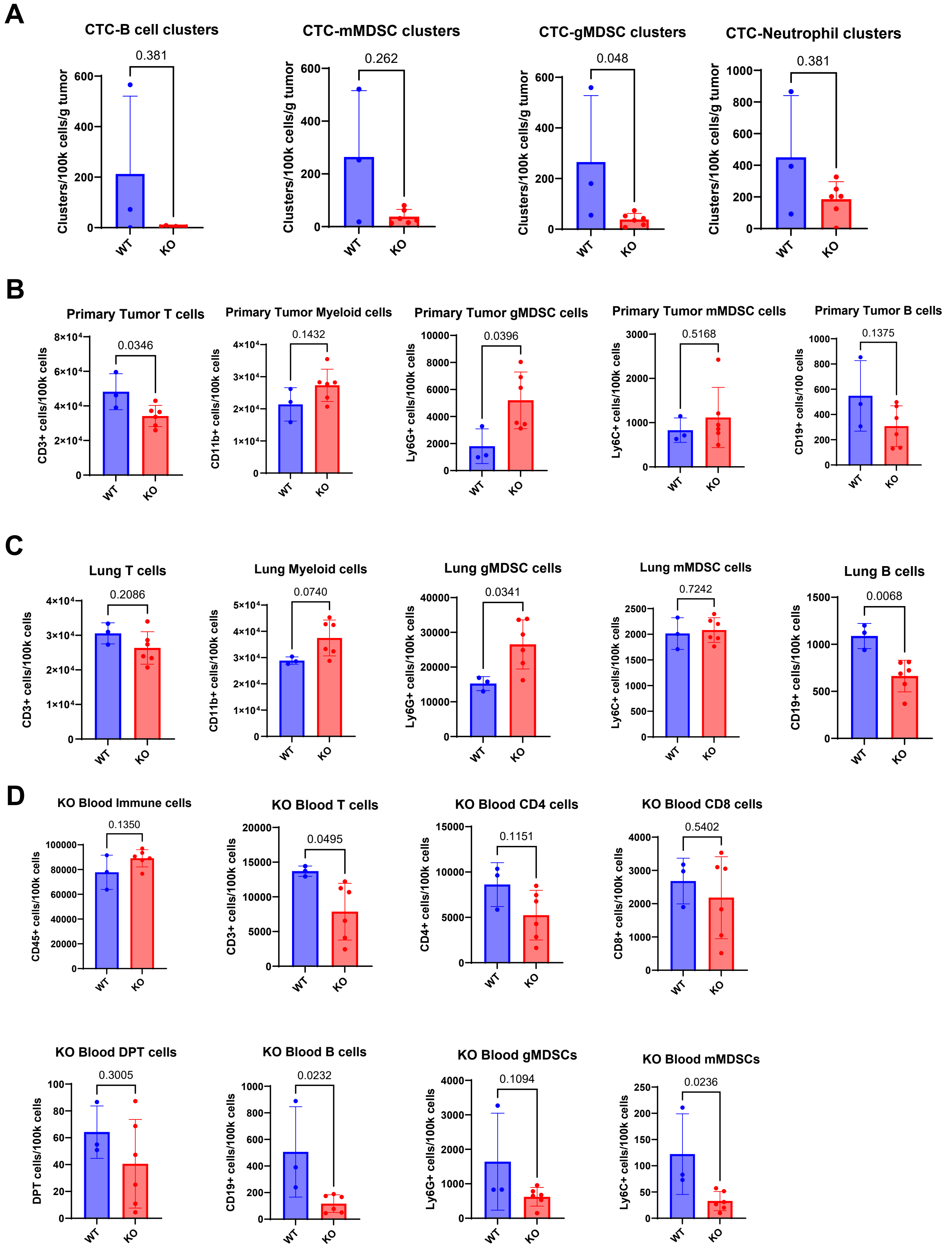


**Figure S9. Immune cell profiles in heterotypic CTC clusters, primary tumors, lungs, and blood.**

**A.** Bar graphs of various blood heterotypic WBC-CTC clusters, including B cells, mMDSC (CD11b+Ly6C+), gMDSC (CD11b+Ly6G+), and neutrophils (CD11b+Ly6G+Ly6C+) in the blood of 4T1 tumor, WT and Vcam1 KO (KO) mice on Day 9 after orthotopic implantation in Fig 5E.

**B-D.** Flow cytometry analysis of T cells ($CD3^{+}$), myeloid cells (CD11b+), gMDSCs (CD11b+Ly6G+), mMDSCs (CD11b+Ly6C+), and B cells (CD19+) in primary tumor (B), lungs (C), and blood (D) of 4T1-WT or 4T1-*Vcam1* KO tumor-bearing mice. Unpaired two-tail t-test, n = 3 for WT control, n = 6 for KO.

### Figure S10

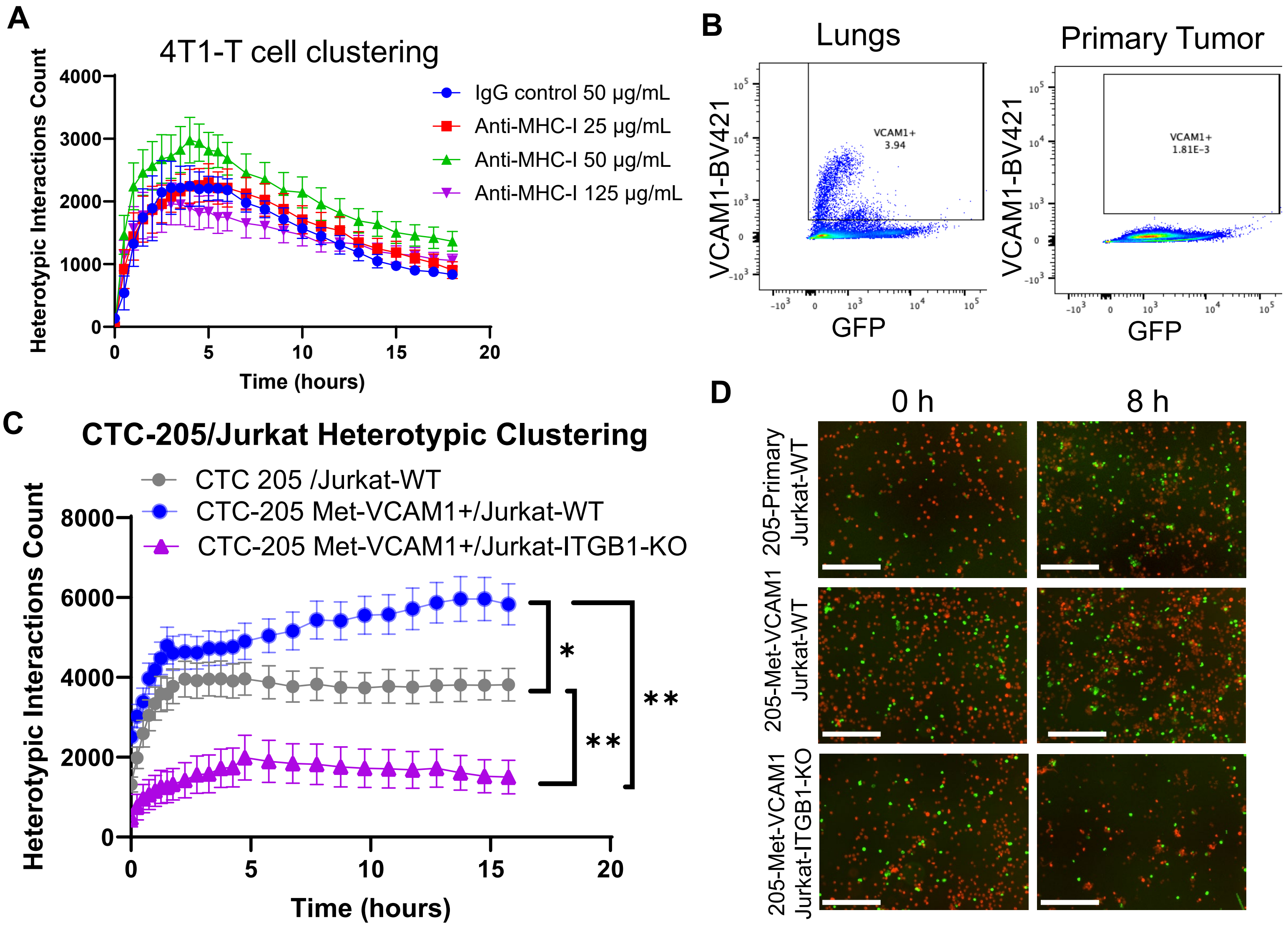


**Figure S10. ITGB1 and VCAM1 as mediators of T cell-tumor cell clustering.**

**A.** Curves depicting heterotypic interactions between mouse splenic T cells and 4T1 cells with pre-treatment of neutralizing control IgG antibody or MHC-I antibody. Graphs represent mean ± standard error of the mean. N = 6 biological replicates.

**B.** Flow cytometry plots showing VCAM1 expression in PDX-CTC-205 lung metastatic lesions (left) or primary tumor (right).

**C-D.** Curves (C) and Representative images (D) depicting heterotypic interactions between Jurkat-Cas9-WT or ITGB1KO cells and PDX-CTC-205 primary tumor cells or lung metastatic lesions. *, $p < 0.05$; **, $p < 0.01$. N = 6 biological replicates. Graphs represent mean ± standard error of the mean.

### Figure S11

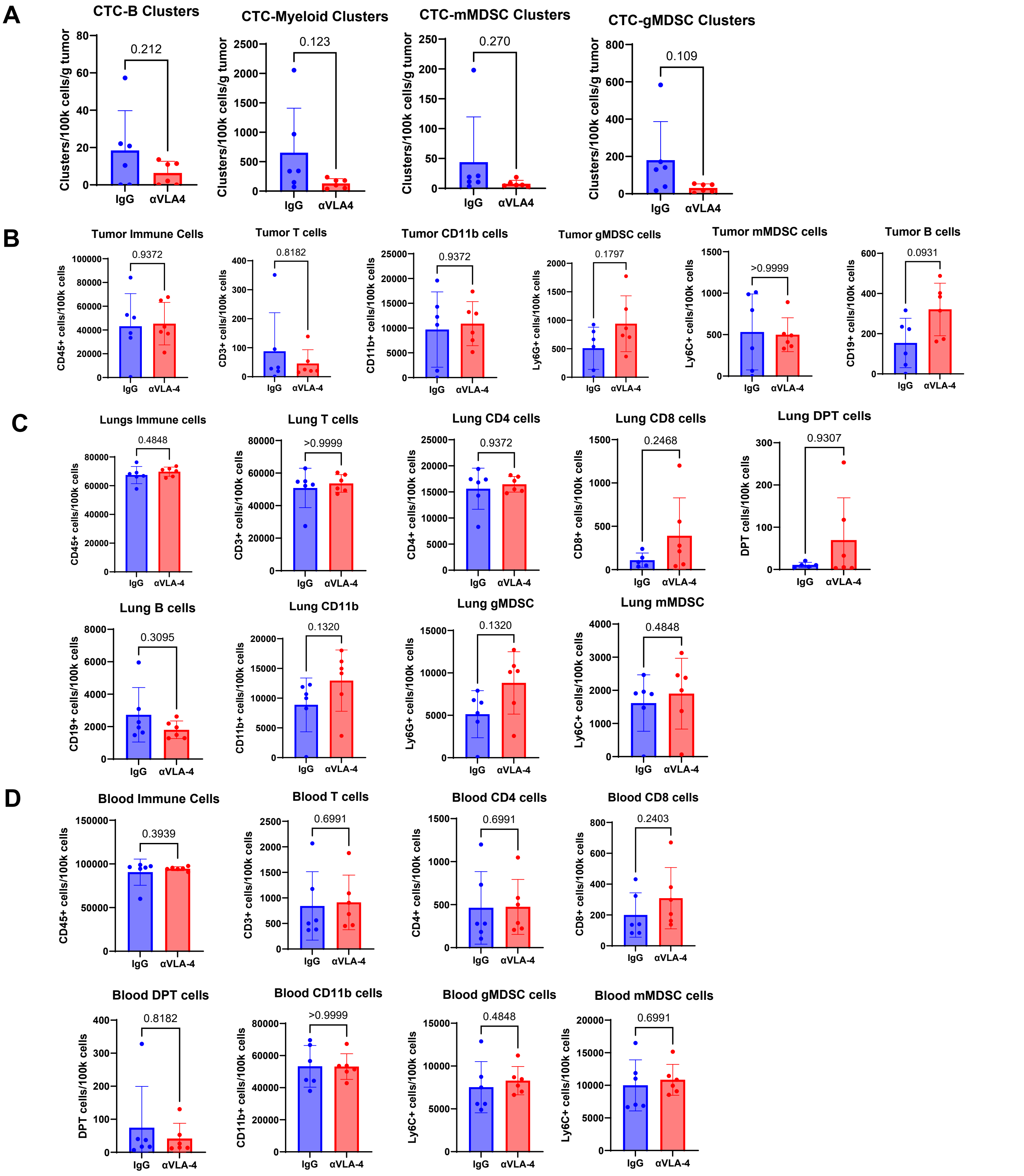


**Figure S11. Immune cell profiles of primary tumors, lungs, and blood cells from the αVLA4 treated mice and the IgG control (related to Figure 5 I-L).**

**A-D.** Bar graphs of flow cytometry data of immune cells ($CD45^+$), T cells ($CD3^+$), myeloid cells ($CD11b^+$), gMDSCs ($CD11b^+Ly6G^+$), mMDSCs ($CD11b^+Ly6C^+$), and B cells ($CD19^+$) in CTC-WBC clusters (A), primary tumor (B), lungs (C), and blood (D) of IgG or αVLA4 (anti-VLA4) treated mice. Unpaired two-tail t-test, N = 6 mice each group.
